# A Glycolysis–Calcineurin Regulatory Axis Orchestrates Titan Cell Formation in *Cryptococcus neoformans*

**DOI:** 10.1101/2025.09.02.673682

**Authors:** Pallavi S Phatak, Sudharsan Mathivathanan, Dhrumi Shah, Ishvarya Suresh, Mary Shejo, Santosh Kumar Das, Sriram Varahan

**Author notes:** Corresponding Author: Dr. Sriram Varahan, Scientist E & Principal Investigator, CSIR-Centre for Cellular and Molecular Biology (CCMB).

## Abstract

*Cryptococcus neoformans* is an opportunistic fungal pathogen that causes pulmonary infections and life-threatening meningoencephalitis in immunocompromised individuals. In addition to the polysaccharide capsule and melanin, titan cell formation is a key virulence trait that promotes immune evasion and disease progression. Despite the established role of titan cells in *C. neoformans* pathogenesis, the molecular mechanisms governing their formation remain poorly understood. Here, we demonstrate that glycolysis is critical for titan cell formation in *C. neoformans*. Pharmacological inhibition or genetic disruption of glycolysis significantly impaired titanization, whereas exogenous cAMP add-back restored the defect. To elucidate the underlying mechanism, we performed comparative RNA-seq analyses of wild- type, *hxk2*Δ, and *hxk2*Δ supplemented with cAMP under titan cell inducing conditions. Transcriptomic analyses revealed significant downregulation of a large subset of calcineurin-responsive genes in the *hxk2*Δ mutant, many of which were restored upon cAMP supplementation. Consistent with these findings, pharmacological inhibition of calcineurin using FK506 or cyclosporin A markedly reduced titan cell formation, establishing an essential role for calcineurin signaling in this morphological transition. Furthermore, supplementation with CaCl_2_ rescued the titanization defect of the *hxk2*Δ mutant, whereas chelation of extracellular calcium with EGTA significantly inhibited titanization in wild-type cells. Using the calcium-sensitive dye, we found that intracellular calcium levels were substantially reduced in the *hxk2*Δ mutant and were restored by CaCl_2_ or cAMP supplementation. Collectively, our findings uncover a previously unrecognized glycolysis–calcineurin regulatory axis that governs titan cell formation and establishes a direct mechanistic link between central carbon metabolism, calcium homeostasis and fungal morphogenesis.

**Summary:** Titan cell formation is a critical virulence trait that promotes immune evasion and disease progression of *Cryptococcus neoformans*. However, the mechanisms governing this morphological transition remain poorly understood. Here, we demonstrate that glycolysis is essential for titan cell formation and uncover a previously unrecognized link between glycolysis, calcium homeostasis, and calcineurin signaling. Comparative transcriptomics, pharmacological perturbation, calcium rescue experiments, and intracellular calcium measurements reveal that disruption of glycolysis attenuates calcineurin signaling by reducing intracellular calcium levels, thereby impairing titanization. These findings establish a novel glycolysis–calcineurin regulatory axis controlling fungal morphogenesis in *C. neoformans*.

## Introduction

*Cryptococcus neoformans* is an opportunistic fungal pathogen that significantly impacts global public health, and causes life-threatening infections especially in immunocompromised individuals (Zhao et al. 2023). This pathogen is responsible for causing cryptococcosis that primarily manifests as a pulmonary infection but can disseminate to cause severe meningoencephalitis (Rajasingham et al. 2017; Chen et al. 2018). With an estimated 220,000 cases of cryptococcal meningitis occurring annually worldwide, resulting in approximately 180,000 deaths, *C. neoformans* has been designated as a critical priority fungal pathogen by the World Health Organization (WHO) (Rajasingham et al. 2017; Zhao et al. 2023). The pathogenicity of *C. neoformans* is attributed to several well-characterized virulence factors that facilitate immune evasion and tissue invasion in the host (Zaragoza 2019). The polysaccharide capsule, composed primarily of glucuronoxylomannan and galactoxylomannan, serves as a major virulence determinant by inhibiting phagocytosis and modulating the host immune response in favour of *C. neoformans* survival (Zaragoza et al. 2009; O’Meara and Andrew Alspaugh 2012). Additionally, melanin production through laccase-mediated oxidation of phenolic compounds, provides protection against oxidative stress and antimicrobial peptides, contributing to *C. neoformans* survival within the host (Nosanchuk and Casadevall 2003; Eisenman and Casadevall 2012). In recent years, morphological plasticity has emerged as another critical virulence mechanism in *C. neoformans* (Okagaki et al. 2010). While the organism typically exists as a budding yeast with cells ranging from 5-7 μm in diameter, several studies have shown that, upon host entry, *C. neoformans* undergoes a dramatic morphological transition resulting in the formation of unusually large cells known as ‘Titan cells’ (Cruickshank et al. 1973; Crabtree et al. 2012; Zhou and Ballou 2018). Titan cells are defined as enlarged cells with a diameter of ≥10 μm (with some cells reaching sizes up to ∼100 μm) (Zhou and Ballou 2018). Several studies have shown that titan cells are resistant to phagocytosis as well as oxidative and nitrosative stresses within the host (Crabtree et al. 2012; Okagaki and Nielsen 2012; García-Rodas et al. 2019). Similarly, in vivo studies suggest that titan cells play a pivotal role in cryptococcal dissemination and disease progression within the host (Okagaki et al. 2010; García-Barbazán et al. 2016). Apart from increased cell size, titan cells exhibit a high degree of polyploidy, a significantly thickened cell wall and capsule with an altered composition (Mukaremera et al. 2018; Zhou and Ballou 2018; García-Rodas et al. 2019). Titan cells also exhibit significant variations in homeostatic cellular processes compared to typical yeast cells, including alterations in cell cycle progression with an extended G2 phase and G2/M arrest (Zaragoza and Nielsen 2013; Altamirano et al. 2021).

Several signaling pathways have been implicated in the regulation of cryptococcal virulence, among which the cyclic AMP-protein kinase A (cAMP-PKA) and calcineurin pathways are two of the best characterized (Mahmood et al. 2024). The cAMP-PKA pathway regulates multiple virulence-associated traits, including capsule formation, melanin biosynthesis, and titan cell development (Alspaugh et al. 1997; D’Souza and Heitman 2001; Pukkila-Worley and Alspaugh 2004; Pukkila-Worley et al. 2005; Caza and Kronstad 2019). Likewise, calcineurin signaling plays a central role in cryptococcal pathogenesis and is required for growth at host physiological temperature, adaptation to elevated CO₂, maintenance of cell wall integrity, and tolerance to environmental stresses encountered during infection (Chow et al. 2017; Yadav and Heitman 2023; Mahmood et al. 2024). Notably, many of the environmental cues that activate calcineurin signaling (including host physiological temperatures and elevated levels of CO₂) closely resemble those used to induce titan cell formation in vitro (Zaragoza and Nielsen 2013; Zhou and Ballou 2018). Despite its established role in virulence, whether calcineurin contributes to titan cell morphogenesis remains unknown.

Recent studies have increasingly highlighted the importance of metabolic pathways in regulating fungal morphological transitions and virulence-associated programs (Hu et al. 2008; Shah et al. 2026). Rather than functioning solely as providers of energy and biosynthetic intermediates, central metabolic pathways including glycolysis have emerged as active regulators of fungal morphogenesis, stress adaptation, and pathogenicity across multiple different species (Dudek et al. 2010; Price et al. 2011; Laurian et al. 2019; Shah et al. 2025). Recently, a series of landmark studies have established that in vitro titanization assays results in the reproducible generation of titan cells that faithfully recapitulate the various biological signatures of titan cells that are generated in vivo (Dambuza et al. 2018; Hommel et al. 2018; Trevijano-Contador et al. 2018). Interestingly, one of these studies showed that several glycolytic genes were upregulated under in vitro titan cell inducing conditions suggesting a possible link between central carbon metabolism and titan cell formation (Trevijano-Contador et al. 2018). However, the functional significance of glycolysis during titanization and the molecular mechanisms linking metabolic state to this morphogenetic transition remain poorly understood. In the present study, we demonstrate that glycolysis is a critical determinant of titan cell formation in *C. neoformans*. Using a combination of comparative transcriptomic analyses, genetic and pharmacological perturbations, calcium rescue experiments, and intracellular calcium measurements, we uncover a previously unrecognized connection between glycolysis, calcium homeostasis, and calcineurin signaling. We show that disruption of glycolysis reduces intracellular calcium levels thereby attenuating the expression of calcineurin-responsive genes, and impairs titan cell formation, whereas exogenous supplementation with calcium restores these defects. Furthermore, pharmacological inhibition of calcineurin phenocopies the glycolytic mutant, establishing calcineurin as a critical downstream regulator of titanization. Remarkably, supplementation with exogenous cAMP, a well-established inducer of titan cell formation (Pukkila-Worley and Alspaugh 2004; Okagaki et al. 2011), restored intracellular calcium levels in the glycolytic mutant to near wild-type levels. Restoration of calcium homeostasis was associated with recovery of the expression of a substantial fraction of calcineurin-responsive genes that were downregulated in the glycolysis mutant, as revealed by comparative transcriptomic analyses. Consistent with these effects, cAMP supplementation also rescued the titan cell formation defect caused by glycolysis perturbation. Collectively, our findings identify a novel glycolysis-calcineurin regulatory axis governing titan cell formation in *C. neoformans* and provide the first evidence that central carbon metabolism directly regulates calcineurin pathway activity in a fungal pathogen.

## Results

### Perturbation of Glycolysis Attenuates Titan Cell Formation in *C. neoformans*

To investigate the functional requirement of glycolysis in titan cell formation, we tested the ability of *C. neoformans* to undergo titanization, using the previously established in vitro titan cell assay (Trevijano-Contador et al. 2018). Briefly, wild-type *C. neoformans* strain H99 was cultured in Titan Cell Medium (TCM) in the presence and absence of sub-inhibitory concentrations of well-established glycolysis inhibitors, 2-Deoxy-D-Glucose (2DG) and sodium citrate (NaCi), which target the early and rate-limiting steps of the glycolysis pathway and titan cell formation was subsequently assessed (Trevijano-Contador et al. 2018) (Fig. 1a). 2DG, a glucose analogue, acts as a competitive inhibitor of hexokinase (Hxk) and phosphoglucose isomerase (Pgi), thereby perturbing glycolysis at its initial steps (Cramer and Woodward 1952). Similarly, NaCi, interferes with the activity of phosphofructokinase (Pfk), which is a key rate-limiting enzyme in the glycolysis pathway (Yoshino and Murakami 1982). Previous studies have shown that when *C. neoformans* is grown in titan cell inducing medium, a heterogeneous population of cells emerges, wherein cell size ranges from 2 µm to ≥10 µm, and cells that are 2–3 µm in diameter have been reported as titanides (Feldmesser et al. 2001; Dambuza et al. 2018). Consistent with the literature, we also observed a heterogeneous population of cells including previously defined titanides (2-3 µm), regular size yeast cells (5- 7 µm), cells between 7 µm to 10 µm and cells ≥10 µm. In our study, we focused on the cells that are ≥10 µm, designated as___titan cells and___the percentage of titan cells were quantified as the ratio of titan cells (≥10 µm) to the total number of cells. Our data clearly demonstrates that sub-inhibitory concentrations of 2DG and NaCi significantly attenuate the ability of *C. neoformans* to undergo titan cell formation (Fig. 1b and c; Supplementary Fig. 1a). Notably, no significant alteration in the titanide population was observed under glycolysis-perturbed conditions following 72 hours of incubation (Supplementary Fig. 1b). To rule out the possibility that the phenotype we observed is due to a general growth defect, we carried out growth curve analysis of *C. neoformans* in TCM, in the presence and absence of glycolysis inhibitors. Our results clearly demonstrate that the reduction in titan cell formation is not due to a general growth defect, as presence of sub-inhibitory concentrations of 2DG and NaCi did not affect the overall growth of the wild-type strain (Fig. 1d). Titan cells are known to have a thicker and more complex capsule structure compared to the capsule of yeast cells. Additionally, in the field, titan cells have been defined by their cell size either including or excluding the capsule (García-Rodas et al. 2019). Increased capsule expansion is thought to be an important morphological hallmark of titan cells, and given this, we wanted to assess whether the perturbation of glycolysis affects titan cell associated capsule thickness, and consequently, overall cell size including the capsule. In order to do this, wild-type cells were grown in TCM in the presence and absence of sub-inhibitory concentrations of glycolysis inhibitors. After incubation, the capsule was visualized using India ink staining and capsule thickness was measured and quantified. Our data shows that the addition of 2DG or NaCi resulted in a significant reduction in overall capsule thickness and total cell diameter (Fig. 1e and f; Supplementary Fig. 1c). This result demonstrates that the perturbation of glycolysis disrupts titan cell associated capsule expansion.

**Fig. 1.**
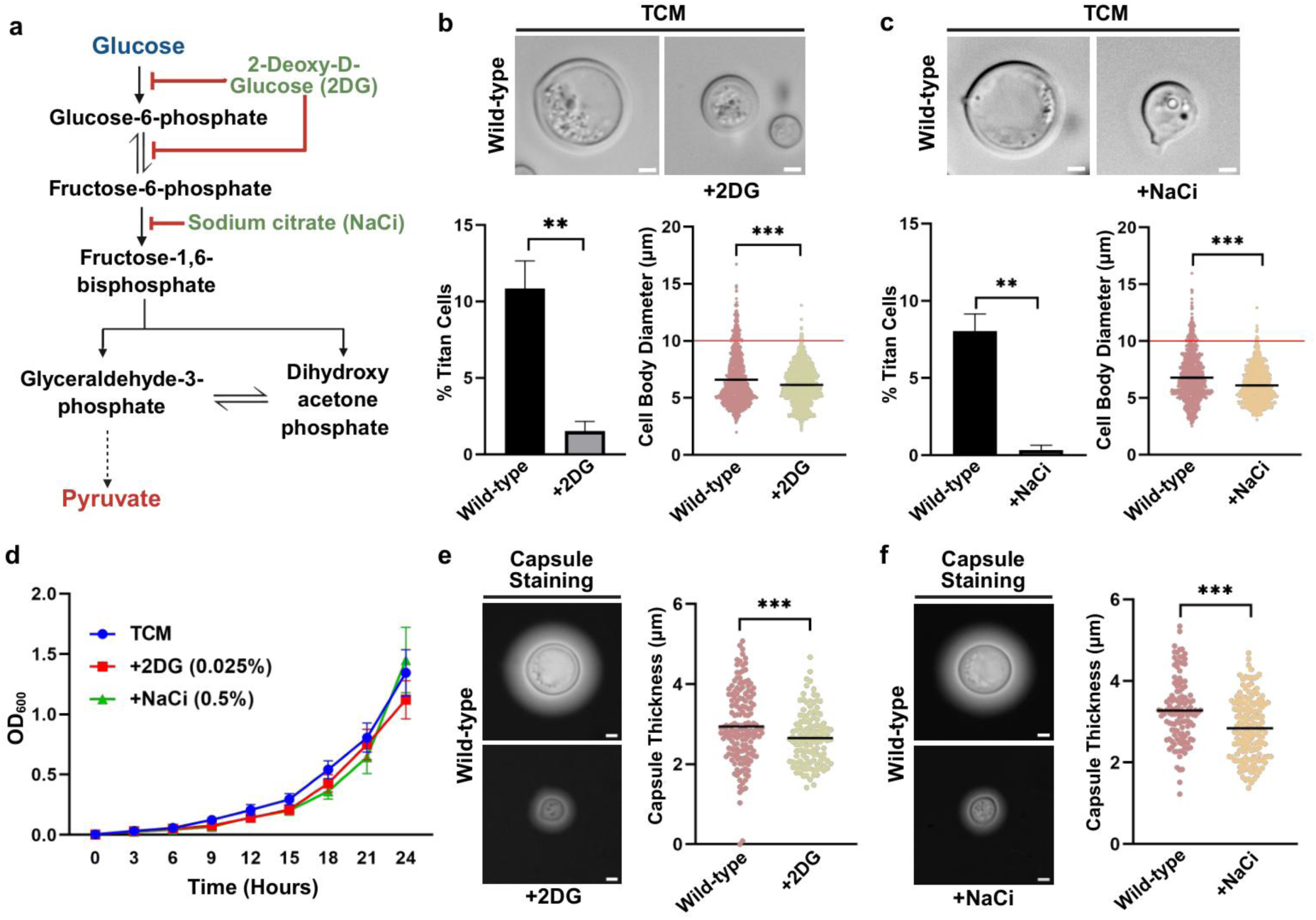
Effect of Glycolysis Inhibitors on Titan Cell Formation in *C. neoformans*. a) Schematic overview of glycolysis and its inhibition by glycolysis inhibitors (2-Deoxy-D-Glucose (2DG) and sodium citrate (NaCi)). b) Wild- type strain was cultured in Titan Cell Medium (TCM) with and without inhibitor (2DG (0.025%)) at 37 °C with 5% CO_2_ for 72 hours. Cells were fixed and imaged using Zeiss Apotome microscope. Percentage of titan cells was quantified as a ratio of cells with a diameter ≥10 µm to the total number of cells. Cell diameter was measured using Zen 2.3 software. Threshold for cell body diameter of titan cells is indicated by the red line. More than 400 cells were counted for each condition. Statistical analysis was performed using unpaired t-test, ***(P<0.001) and **(P<0.01). Error bars represent SEM. Scale bars represent 2 µm. c) Wild-type strain was cultured in TCM with and without inhibitor (NaCi (0.5%)) at 37 °C with 5% CO_2_ for 72 hours. Cells were fixed and imaged using Zeiss Apotome microscope. Percentage of titan cells was quantified as a ratio of cells with a diameter ≥10 µm to the total number of cells. Cell diameter was measured using Zen 2.3 software. Threshold for cell body diameter of titan cells is indicated by the red line. More than 400 cells were counted for each condition. Statistical analysis was performed using unpaired t-test, ***(P<0.001) and **(P<0.01). Error bars represent SEM. Scale bars represents 2 µm. d) Growth curve was performed to monitor overall growth of wild-type strain under titan cell inducing conditions in the presence and absence of glycolysis inhibitors (2DG or NaCi). Overnight grown culture of wild-type strain (H99) was diluted to OD_600_=0.01 in fresh TCM medium with and without 2DG (0.025%) or NaCi (0.5%) and allowed to grow at 37 °C for 24 hours. OD_600_ was recorded at 3-hour intervals. e) For capsule visualization, wild-type strain was cultured in TCM with and without 2DG at 37 °C with 5% CO_2_ for 72 hours, cells were stained with India ink and imaging was done using Zeiss Apotome microscope. Capsule thickness was measured using Zen 2.3 software. Approximately 150 cells were counted for each condition. Statistical analysis was performed using unpaired t-test, ***(P<0.001). Error bars represent SEM Scale bars represent 2 µm. f) For capsule visualization, wild-type strain was cultured in TCM with and without NaCi at 37 °C with 5% CO_2_ for 72 hours, cells were stained with India ink and imaging was done using Zeiss Apotome microscope. Capsule thickness was measured using Zen 2.3 software. Approximately 150 cells were counted for each condition. Statistical analysis was performed using unpaired t-test; ***(P<0.001). Error bars represent SEM. Scale bars represent 2 µm. This figure was created using BioRender. https://BioRender.com/ny8fork.

Given that, 2DG and NaCi exhibited a strong inhibitory effect on titan cell formation, we wanted to corroborate these observations by using pertinent genetic knockout strains that are compromised in their ability to metabolize glucose via glycolysis (Price et al. 2011). In order to do this, we tested the ability of knockout strains which lack the genes that encode for enzymes that are involved in the glycolysis pathway including, Hxk2 (Hexokinase 2) and Pyk1 (Pyruvate kinase 1) (Price et al. 2011) (Fig. 2a), to undergo titanization. In *C. neoformans*, *HXK2* encodes for the enzyme that catalyses the phosphorylation of glucose to glucose-6- phosphate (Idnurm et al. 2007), while *PYK1* encodes for the enzyme responsible for converting phosphoenolpyruvate to pyruvate (Price et al. 2011). Briefly, *hxk2*Δ and *pyk1*Δ (*hxk2*Δ refers to the *HXK2* knockout strain and *pyk1*Δ refers to the *PYK1* knockout strain) along with the wild-type were cultured in titan cell inducing conditions. Our result demonstrates that the deletion of *HXK2* and *PYK1* resulted in a significant reduction in titan cell formation, compared to the wild-type strain (Fig. 2b and c; Supplementary Fig. 2a). Similar to our observation with glycolysis inhibitors, no significant alteration in the titanide population was observed in glycolysis-deficient strains following 72 hours of incubation (Supplementary Fig. 3a and b). To rule out the possibility that the phenotype we observed is due to a general growth defect, we performed growth curve analysis of the wild-type and glycolysis mutants (*hxk2*Δ and *pyk1*Δ) in TCM. Our data suggests that the reduction in titan cell formation is not due to general growth defects as the mutants were able to grow similar to the wild-type strain in TCM (Fig. 2d). Given that the perturbation of glycolysis using 2DG or NaCi leads to reduced titan cell associated capsule thickness in the wild-type, we wanted to corroborate these observations by using pertinent genetic knockout strains (*hxk2*Δ and *pyk1*Δ) that are compromised in their ability to undergo glycolysis. In order to do this, glycolysis knockout strains including*, hxk2*Δ and *pyk1*Δ along with the wild-type were cultured in TCM. After incubation, the capsule was visualized using India ink staining and capsule thickness was measured and quantified. Bright field microscopy revealed that glycolysis knockout strains exhibited a significant decrease in capsule thickness compared to the wild-type strain (Fig. 2e and f; Supplementary Fig. 2b).

**Fig. 2.**
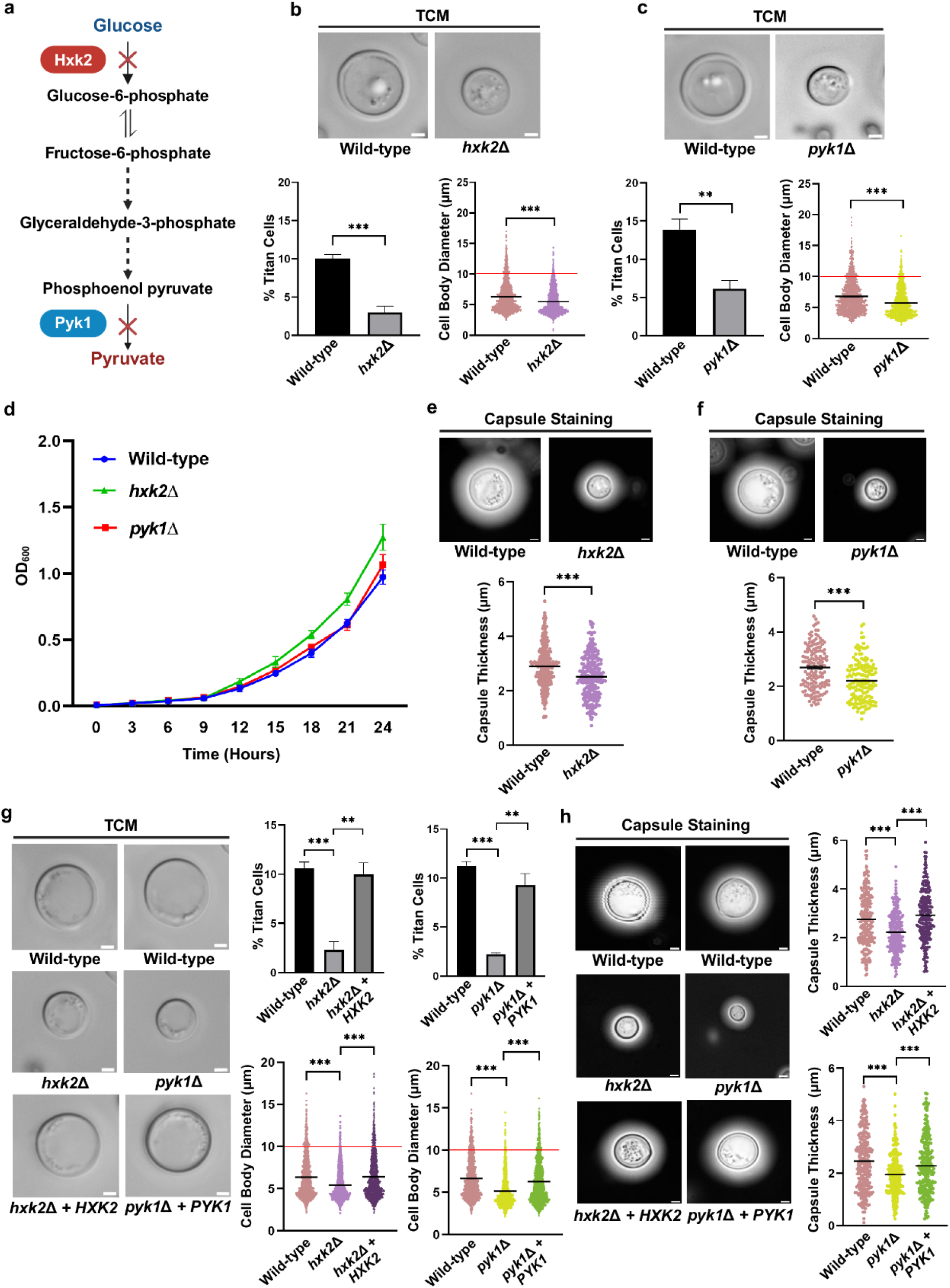
Genetic Perturbation of Glycolysis Attenuates Titan Cell Formation in *C. neoformans*. a) Overview of glycolysis pathway showing targeted gene deletions. b) Wild-type and *hxk2*Δ strains were cultured in Titan Cell Medium (TCM) at 37 °C with 5% CO_2_ for 72 hours. Cells were fixed and imaged using Zeiss Apotome microscope. Percentage of titan cells was quantified as a ratio of cells with a diameter ≥10 µm to total number of cells. Cell diameter was measured using Zen 2.3 software. Threshold for cell body diameter of titan cells is indicated by the red line. More than 400 cells were counted for each condition. Statistical analysis was performed using unpaired t- test; ***(P<0.001). Error bars represent SEM. Scale bars represent 2 µm. c) Wild-type and *pyk1Δ* strains were cultured in TCM at 37 °C with 5% CO_2_ for 72 hours. Cells were fixed and imaged using Zeiss Apotome microscope. Percentage of titan cells was quantified as a ratio of cells with a diameter ≥10 µm to total number of cells. Cell diameter was measured using Zen 2.3 software. Threshold for cell body diameter of titan cells is indicated by the red line. More than 400 cells were counted for each condition. Statistical analysis was performed using unpaired t- test, ***(P<0.001) and **(P<0.01). Error bars represent SEM. Scale bars represent 2 µm. d) Growth curve was performed to monitor overall growth of wild-type and glycolysis mutants (*hxk2*Δ and *pyk1*Δ) under titan cell inducing conditions. Overnight grown culture of wild-type strain (H99) and glycolysis mutants (*hxk2*Δ and *pyk1*Δ) were diluted to OD_600_=0.01 in fresh TCM medium and allowed to grow at 37 °C for 24 hours. OD_600_ was recorded at 3- hour intervals. e) For capsule visualization, wild-type, and *hxk2*Δ strains were cultured in TCM at 37 °C with 5% CO_2_ for 72 hours, cells were stained with India ink and imaging was done using Zeiss Apotome microscope. Capsule thickness was measured using Zen 2.3 software. Approximately 150 cells were counted for each condition. Statistical analysis was performed using unpaired t-test, ***(P<0.001). Error bars represent SEM. Scale bars represent 2 µm. f) For capsule visualization, wild-type and *pyk1*Δ strains were cultured in TCM at 37 °C with 5% CO_2_ for 72 hours, cells were stained with India ink and imaging was done using Zeiss Apotome microscope. Capsule thickness was measured using Zen 2.3 software. Approximately 150 cells were counted for each condition. Statistical analysis was performed using unpaired t-test, ***(P<0.001). Error bars represent SEM. Scale bars represent 2 µm. g) Wild-type, *hxk2*Δ*, pyk1*Δ*, hxk2*Δ *+ HXK2* and *pyk1*Δ *+ PYK1* strains were cultured in TCM at 37 °C with 5% CO_2_ for 72 hours. Cells were fixed and imaged using Zeiss Apotome microscope. Percentage of titan cells was quantified as a ratio of cells with a diameter ≥10 µm to total number of cells. Cell diameter was measured using Zen 2.3 software. Threshold for cell body diameter of titan cells is indicated by the red line. More than 400 cells were counted for each condition. Statistical analysis was performed using unpaired t-test, ***(P<0.001) and **(P<0.01). Error bars represent SEM. Scale bars represent 2 µm. h) For capsule visualization, wild-type, *hxk2*Δ*, pyk1*Δ*, hxk2*Δ *+ HXK2* and *pyk1*Δ *+ PYK1* strains were cultured in TCM at 37 °C with 5% CO_2_ for 72 hours, cells were stained with India Ink and imaging was done using Zeiss Apotome microscope. Capsule thickness was measured using Zen 2.3 software. Approximately 150 cells were counted for each condition. Statistical analysis was performed using unpaired t-test; ***(P<0.001). Error bars represent SEM. Scale bars represent 2 µm. This figure was created using BioRender. https://BioRender.com/izvfw77.

To confirm that the defects observed in titan cell formation, in the glycolysis mutants (*hxk2*Δ and *pyk1*Δ), were not due to any off-target integrations, chromosomal deletions/duplications, and tandem integrations caused by the transformation procedure, we validated our results using complemented strains for these mutants. Our results demonstrated that the complemented strains, *hxk2*Δ + *HXK2* and *pyk1*Δ + *PYK1* exhibited complete rescue of the titanization defects exhibited by the mutants, *hxk2*Δ and *pyk1*Δ (Fig. 2g; Supplementary Fig. 2c). These complementation strains were also tested for titan cell associated capsule formation and total cell size, using India ink staining. As shown in the Fig. 2h, complemented strains *hxk2*Δ + *HXK2* and *pyk1*Δ + *PYK1* exhibited complete rescue of the defects observed in titan cell associated capsule formation in the glycolysis mutants *hxk2*Δ and *pyk1*Δ (Supplementary Fig. 2d). Taken together, our results clearly demonstrate that the ability of *C. neoformans* to efficiently metabolize glucose via glycolysis is critical for titan cell formation.

### Perturbation of Glycolysis Attenuates Calcineurin-mediated Transcriptional Regulation under Titan Cell Inducing Conditions in *C. neoformans*

Our previous results demonstrated that perturbation of glycolysis, whether through pharmacological inhibition or the use of glycolysis-deficient mutant strains, leads to a significant reduction in titan cell formation in *C. neoformans*. Disruption of glycolysis pathway not only alters the overall metabolic state of the cell but also influences a broad spectrum of downstream processes, including cell wall biosynthesis and cell cycle progression (Kalucka et al. 2015; Karri et al. 2024; Kierans and Taylor 2024). Beyond its role in energy generation, several glycolytic intermediates function as second messengers involved in various key signaling pathways across different species. In fungal systems, for example, fructose-1,6- bisphosphate (Fru-1,6-BP) has been shown to activate the cAMP-PKA pathway by directly stimulating the guanine nucleotide exchange factor Cdc25, which in turn activates the GTPase Ras, thereby coupling glycolytic flux to downstream cAMP signaling (Peeters et al. 2017). Additionally, diacylglycerol (DAG), a lipid second messenger biosynthetically linked to the glycolytic intermediate dihydroxyacetone phosphate (DHAP) via the glycerophospholipid synthesis pathway, is a well-established activator of Protein Kinase C (PKC), which governs the Cell Wall Integrity (CWI) signaling cascade in fungi (Athenstaedt et al. 1999; Levin 2005; Gerik et al. 2008; Heinisch and Rodicio 2018; Athenstaedt 2021). Importantly, in *C. neoformans*, several of these signaling networks, including the cAMP-PKA and CWI pathways, have been implicated in the regulation of titan cell formation (Pukkila-Worley and Alspaugh 2004; Trevijano-Contador et al. 2018; de Oliveira et al. 2021; Mahmood et al. 2024). To gain mechanistic into how glycolysis regulates titan cell formation, we performed comparative RNA-seq analysis on wild-type and glycolysis mutant strain grown under titan cell inducing conditions (Fig. 3a). While in vivo and in vitro studies indicate that titan cell populations were maximum at approximately 72 hours (Dambuza et al. 2018; Trevijano-Contador et al. 2018), the regulatory signaling and gene expression changes driving this morphogenesis likely occur much earlier. Consequently, while we maintained a 72 hours incubation to confirm robust titanization in our in vitro assays, we selected the 24-hour time point for transcriptomic analysis to gain mechanistic insights into how glycolysis regulates this morphological transition. Briefly, *hxk2*Δ mutant along with wild-type strain were cultured in titan cell inducing conditions for 24 hours and RNA was extracted using hot-phenol method. Cell lysis was done by beat-beating as described (Material and Methods).

**Fig. 3.**
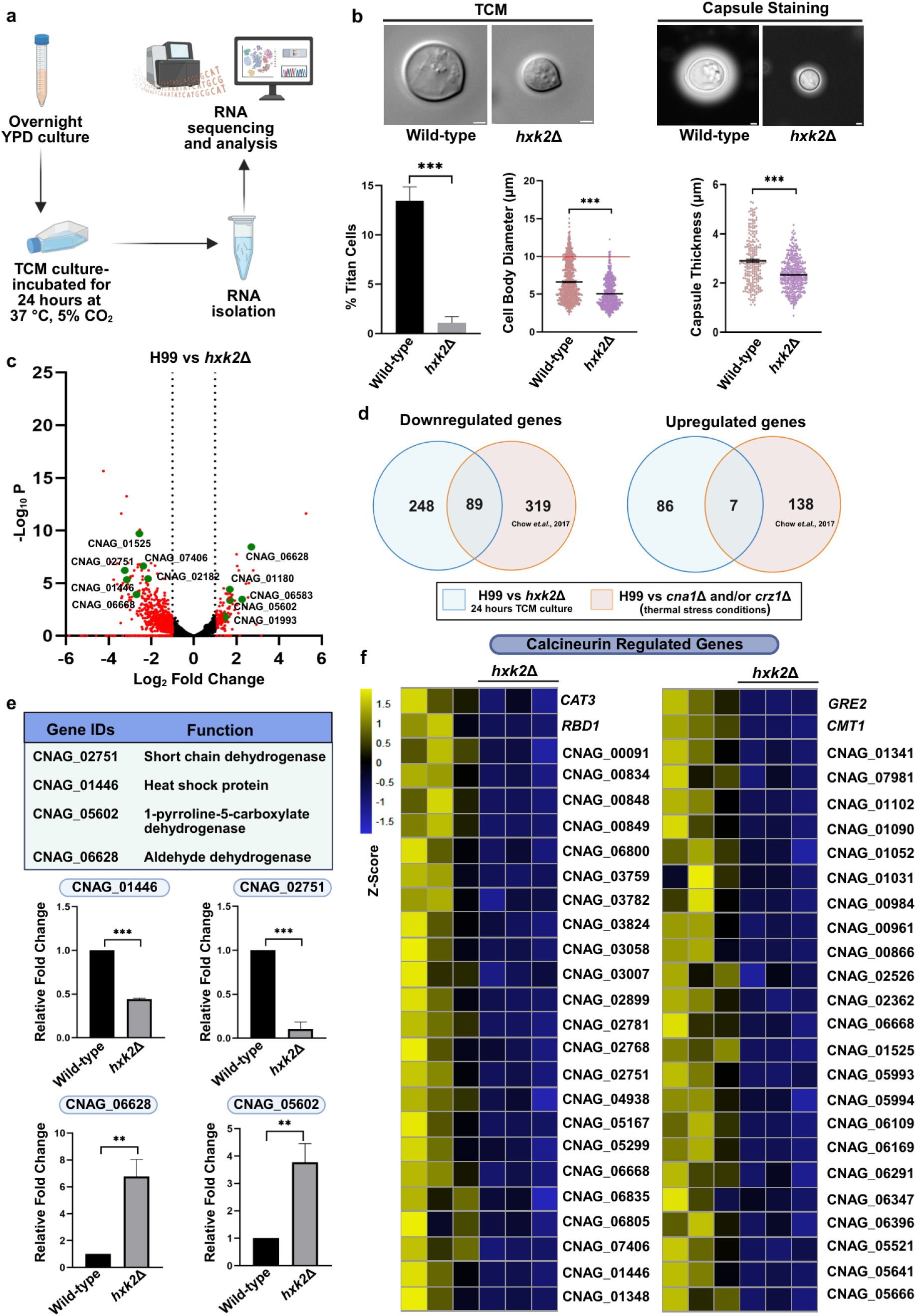
Perturbation of Glycolysis Attenuates Calcineurin-mediated Transcriptional Regulation under Titan Cell Inducing Conditions in *C. neoformans*. a) Schematic overview of steps involved in RNA isolation and sequencing. b) Wild-type and *hxk2*Δ strains were cultured in TCM at 37 °C with 5% CO_2_ for 24 hours. Cells were fixed and imaged using Zeiss Apotome microscope. Percentage of titan cells was quantified as a ratio of cells with a diameter ≥10 µm to total number of cells. Cell diameter was measured using Zen 2.3 software. Threshold for cell body diameter of titan cells is indicated by the red line. More than 400 cells were counted for each condition. Statistical analysis was performed using unpaired t-test; ***(P<0.001). Error bars represent SEM. Scale bars represent 2 µm. For capsule visualization cells were stained with India ink and imaging was done using Zeiss Apotome microscope. Capsule thickness was measured using Zen 2.3 software. Approximately 150 cells were counted for each condition. Statistical analysis was performed using unpaired t-test; ***(P<0.001). Error bars represent SEM. Scale bars represent 2 µm. c) Volcano plots represent differentially expressed genes in *hxk*2Δ strain compared to wild-type strain grown under titan cell inducing conditions for 24 hours. Genes which are upregulated (Log_2_ fold change ≥ 1) and downregulated (Log_2_ fold change ≤ -1) are highlighted in red. Some of the significantly upregulated and downregulated genes involved in the calcineurin pathway are highlighted in green. d) Venn diagram representing the overlap between our RNA-seq dataset (wild-type vs *hxk2*Δ under titan cell inducing condition) and dataset published by Chow et al. 2017 (wild-type vs calcineurin mutants under thermal stress condition). e) Validation of RNA-seq data by RT-qPCR analysis for a subset of downregulated genes (CNAG_01446 and CNAG_02751) and upregulated genes (CNAG_06628 and CNAG_05602) in *hxk2*Δ grown under titan cell inducing conditions for 24 hours. Statistical analysis was performed using unpaired t-test, ***(P<0.001) and **(P<0.01). Error bars represent SEM. f) Heatmaps represent a subset of downregulated genes regulated by the calcineurin pathway, in a *hxk2*Δ mutant compared to wild-type strain grown under titan cell inducing conditions. Scale bar represents Z-score. This figure was created using BioRender. https://BioRender.com/ummpg7p.

Before proceeding with the transcriptomic analysis, we monitored the ability of wild-type and *hxk2*Δ mutant to undergo titanization after 24 hours in TCM. As shown in the figure (Fig. 3b; Supplementary Fig. 4a and b), the *hxk2*Δ mutant displayed a clear reduction in titan cell formation compared to the wild-type, and this was further supported by capsule analysis, which was consistent with the observed titanization phenotypes (Fig. 3b; Supplementary Fig. 4a and b). Having validated the phenotypes, we then compared the transcriptomes of wild- type and *hxk2*Δ strains grown under titan cell inducing conditions. This analysis revealed that 430 genes were differentially expressed in the *hxk2*Δ mutant, with the majority being downregulated (337 genes) and a smaller subset being upregulated (93 genes). Fig. 3C represents a volcano plot for the differentially expressed genes in *hxk2*Δ relative to the wild-type strain. Although many of these genes remain poorly characterized in *C. neoformans*, functional categorization of the downregulated genes highlighted the involvement of these in several biologically relevant processes, including cell cycle regulation, cell wall integrity and homeostasis, cellular transport, and cellular response to heat stress (Table 1) (Chow et al. 2017; Yu et al. 2021; Martinez Barrera et al. 2024). In pathogenic fungi such as *C. albicans* and *C. neoformans*, the calcineurin pathway is a key regulator of the cellular heat stress response and is involved in the maintenance of cell wall integrity and homeostasis (Yadav and Heitman 2023). Interestingly, when we compared our list of downregulated genes with the RNA-seq dataset published by Chow et al. (Chow et al. 2017), which identified around 408 genes downregulated in calcineurin-deficient mutants under thermal stress, we found a substantial overlap of 89 genes (Fig. 3d). To visually highlight this overlap, a subset of these common genes was highlighted in green on the volcano plot (Fig. 3c), while the expression patterns of selected genes were further visualized using heat maps (Fig. 3f). This overlap strongly suggests that disruption of glycolysis in the *hxk2*Δ mutant attenuates calcineurin signaling under titan cell inducing conditions, pointing to a previously uncharacterized connection between glycolytic flux and calcineurin-mediated signaling.

**Table 1:**
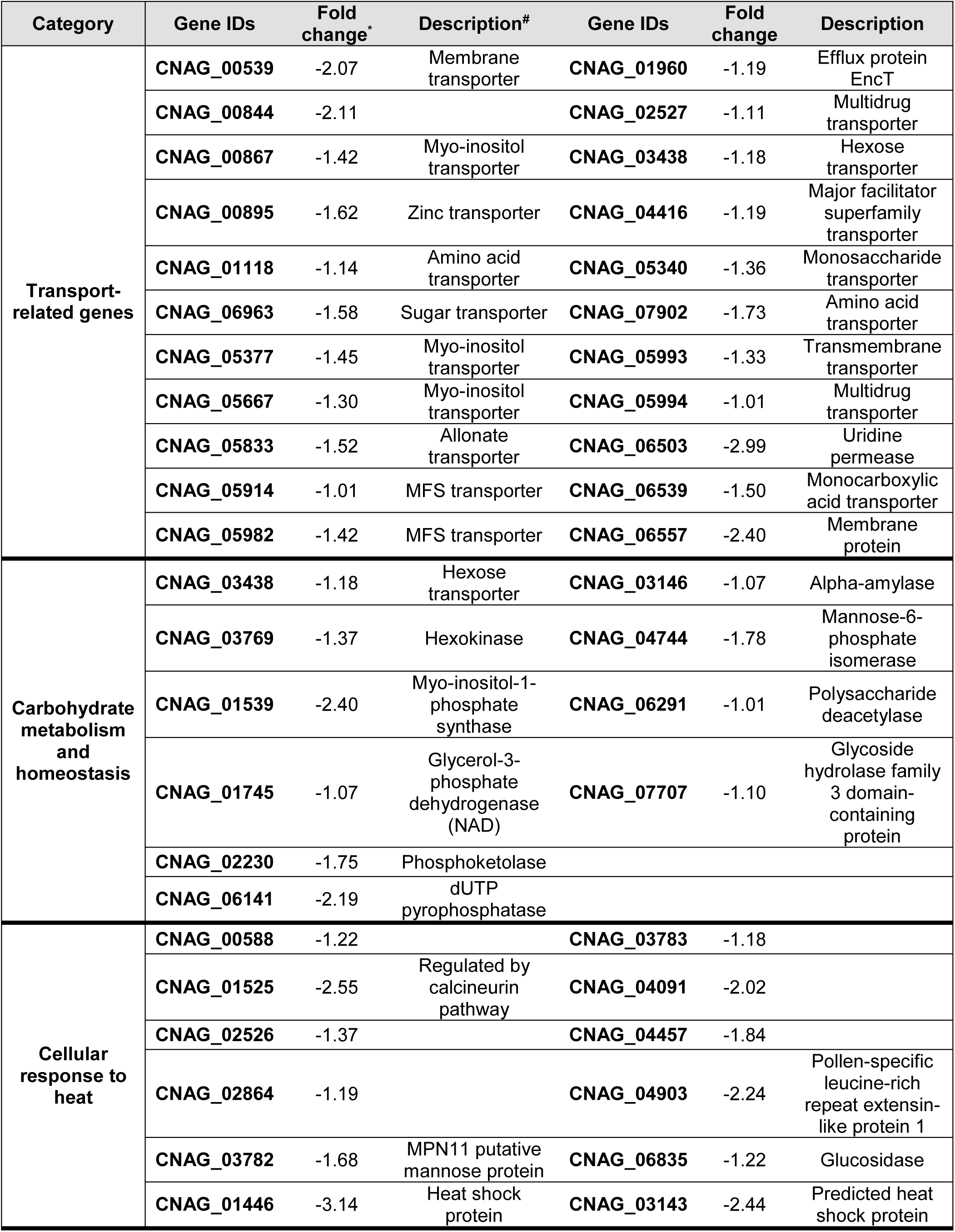
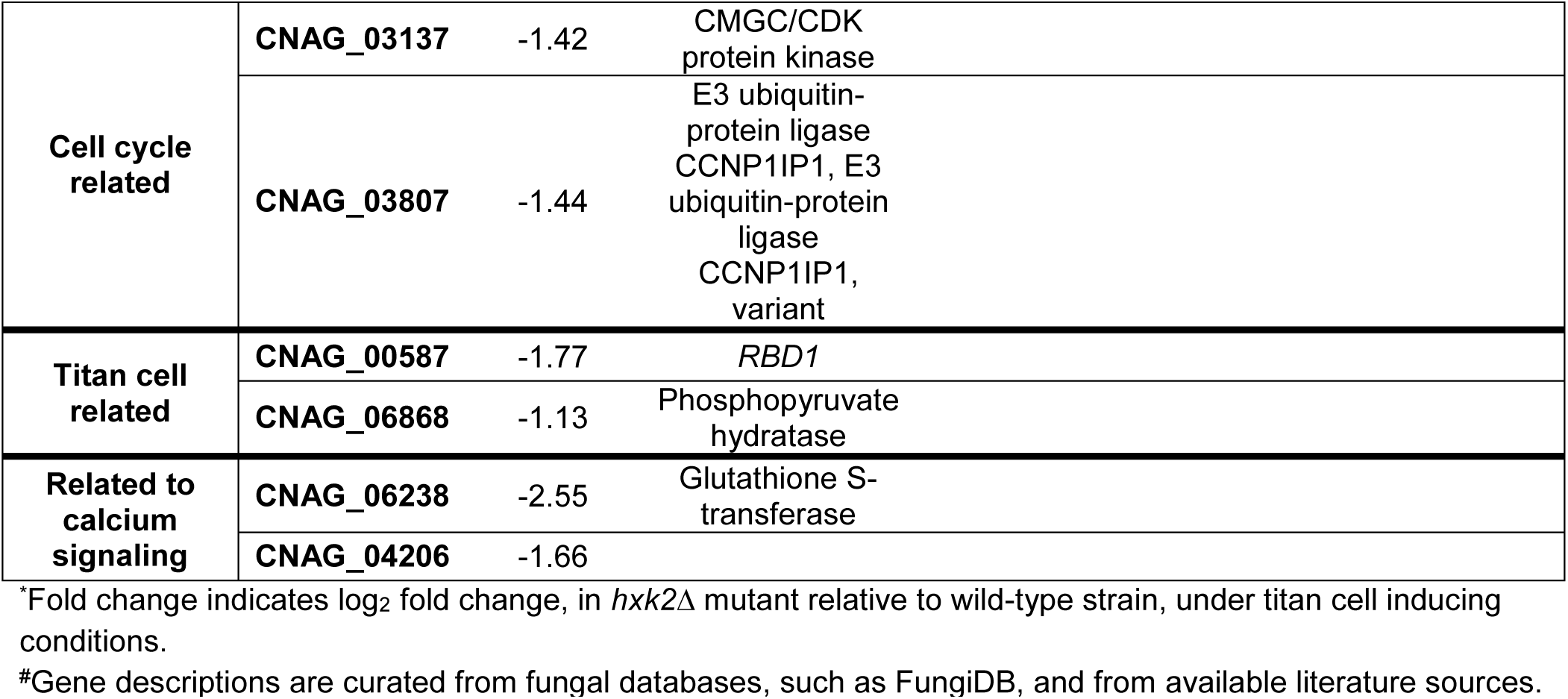
List of Functional Categories of Downregulated Genes in the Glycolysis- Deficient Mutant.

| Category | Gene IDs | Fold change* | Description<sup>#</sup> | Gene IDs | Fold change | Description |
| --- | --- | --- | --- | --- | --- | --- |
| Transport-related genes | CNAG_00539 | -2.07 | Membrane transporter | CNAG_01960 | -1.19 | Efflux protein EncT |
|  | CNAG_00844 | -2.11 |  | CNAG_02527 | -1.11 | Multidrug transporter |
|  | CNAG_00867 | -1.42 | Myo-inositol transporter | CNAG_03438 | -1.18 | Hexose transporter |
|  | CNAG_00895 | -1.62 | Zinc transporter | CNAG_04416 | -1.19 | Major facilitator superfamily transporter |
|  | CNAG_01118 | -1.14 | Amino acid transporter | CNAG_05340 | -1.36 | Monosaccharide transporter |
|  | CNAG_06963 | -1.58 | Sugar transporter | CNAG_07902 | -1.73 | Amino acid transporter |
|  | CNAG_05377 | -1.45 | Myo-inositol transporter | CNAG_05993 | -1.33 | Transmembrane transporter |
|  | CNAG_05667 | -1.30 | Myo-inositol transporter | CNAG_05994 | -1.01 | Multidrug transporter |
|  | CNAG_05833 | -1.52 | Allonate transporter | CNAG_06503 | -2.99 | Uridine permease |
|  | CNAG_05914 | -1.01 | MFS transporter | CNAG_06539 | -1.50 | Monocarboxylic acid transporter |
|  | CNAG_05982 | -1.42 | MFS transporter | CNAG_06557 | -2.40 | Membrane protein |
| Carbohydrate metabolism and homeostasis | CNAG_03438 | -1.18 | Hexose transporter | CNAG_03146 | -1.07 | Alpha-amylase |
|  | CNAG_03769 | -1.37 | Hexokinase | CNAG_04744 | -1.78 | Mannose-6-phosphate isomerase |
|  | CNAG_01539 | -2.40 | Myo-inositol-1-phosphate synthase | CNAG_06291 | -1.01 | Polysaccharide deacetylase |
|  | CNAG_01745 | -1.07 | Glycerol-3-phosphate dehydrogenase (NAD) | CNAG_07707 | -1.10 | Glycoside hydrolase family 3 domain-containing protein |
|  | CNAG_02230 | -1.75 | Phosphoketolase |  |  |  |
|  | CNAG_06141 | -2.19 | dUTP pyrophosphatase |  |  |  |
| Cellular response to heat | CNAG_00588 | -1.22 |  | CNAG_03783 | -1.18 |  |
|  | CNAG_01525 | -2.55 | Regulated by calcineurin pathway | CNAG_04091 | -2.02 |  |
|  | CNAG_02526 | -1.37 |  | CNAG_04457 | -1.84 |  |
|  | CNAG_02864 | -1.19 |  | CNAG_04903 | -2.24 | Pollen-specific leucine-rich repeat extensin-like protein 1 |
|  | CNAG_03782 | -1.68 | MPN11 putative mannose protein | CNAG_06835 | -1.22 | Glucosidase |
|  | CNAG_01446 | -3.14 | Heat shock protein | CNAG_03143 | -2.44 | Predicted heat shock protein |

|  |  |  |  |
| --- | --- | --- | --- |
| Cell cycle related | CNAG_03137 | -1.42 | CMGC/CDK protein kinase |
|  | CNAG_03807 | -1.44 | E3 ubiquitin-protein ligase CCNP1IP1, E3 ubiquitin-protein ligase CCNP1IP1, variant |
| Titan cell related | CNAG_00587 | -1.77 | *RBD1* |
|  | CNAG_06868 | -1.13 | Phosphopyruvate hydratase |
| Related to calcium signaling | CNAG_06238 | -2.55 | Glutathione S-transferase |
|  | CNAG_04206 | -1.66 |  |

\*Fold change indicates log<sub>2</sub> fold change, in *hvk2Δ* mutant relative to wild-type strain, under titan cell inducing conditions.
#Gene descriptions are curated from fungal databases, such as FungiDB, and from available literature sources.

To validate our transcriptomic findings, we performed quantitative real-time PCR (RT-qPCR) to measure the expression of 4 candidate genes representing both significantly downregulated and upregulated genes (These 4 genes are commonly downregulated and upregulated in our RNA-seq dataset as well as in the Chow et al. 2017 study). The RT-qPCR results were largely consistent with our RNA-seq data, confirming the reliability of our transcriptomic analysis and further supporting the observation that glycolysis perturbation influences calcineurin-mediated gene expression in *C. neoformans* (Fig. 3e).

### Inhibition of Calcineurin Impairs Titan Cell Formation in *C. neoformans*

Our previous results clearly establish that functional glycolysis acts as a positive regulator of titan cell formation. To understand the mechanistic basis of this regulation, we performed a comprehensive transcriptomic analysis between the *hxk2*Δ mutant and wild-type *C. neoformans* grown under titan cell inducing conditions. Notably, this analysis revealed that perturbation of glycolysis was associated with a broad suppression of calcineurin-dependent gene expression, suggesting that the two may be functionally connected in the context of titan cell formation. Calcineurin is a calcium-dependent signaling cascade that is broadly conserved across diverse biological systems (Park et al. 2019; Yadav and Heitman 2023). In *C. neoformans* specifically, the calcineurin pathway has been shown to be essential for survival under host-relevant conditions, including the physiological temperature of 37 °C and the high CO₂ environment encountered within the host (Chow et al. 2017; Squizani et al. 2020; Squizani et al. 2021; Yadav and Heitman 2023). Importantly, calcineurin serves as a central regulator of the cellular response to a range of stress conditions, including heat, pH fluctuations, and elevated CO₂ levels (Chadwick et al. 2022; Mahmood et al. 2024). While the literature has established the importance of this pathway in the context of thermotolerance and virulence, its role in capsule formation appears to be more context-dependent (Kmetzsch et al. 2010; Squizani et al. 2020). Strikingly, despite its importance in *C. neoformans* physiology and virulence, the role of the calcineurin pathway in regulating titan cell formation has remained largely unexplored.

We therefore asked whether calcineurin activity is required for titanization. To address this, we treated *C. neoformans* with two well-established pharmacological inhibitors of the calcineurin pathway, Cyclosporin A (CsA) and FK506, and assessed their effect on titan cell formation (Hemenway and Heitman 1999; Chow et al. 2017). Both compounds are widely used as antifungal and immunosuppressive agents, and while they share a common target, they act through distinct mechanisms. CsA binds to cyclophilin A, while FK506 binds to FKBP12. In each case, the resulting drug-protein complex binds to and blocks calcineurin, preventing it from interacting with its substrates and thereby inhibiting downstream signaling (Fig. 4a) (Hemenway and Heitman 1999; Lev et al. 2012; Chow et al. 2017). Importantly, both inhibitors have been previously employed at sub-inhibitory concentrations in fungal studies to selectively perturb calcineurin pathway activity without broadly compromising cellular growth (Lev et al. 2012; Chow et al. 2017; Kalem et al. 2021) . To assess the role of the calcineurin pathway in titan cell formation, we treated wild-type *C. neoformans* with sub-inhibitory concentrations of each inhibitor. Briefly, wild-type *C. neoformans* strain was cultured in titan cell inducing conditions, in the presence and absence of sub-inhibitory concentrations of CsA or FK506. CsA as well as FK506 inhibited titan cell formation significantly, at the tested concentrations (Fig. 4b and C; Supplementary Fig. 5a). To rule out the possibility that the phenotype we observed is due to a general growth defect we performed the growth curve assay in the presence and absence of the lower sub-inhibitory concentrations of CsA and FK506. Our results clearly demonstrate that the reduction in titan cell formation is not due to a general growth defect as presence of sub-inhibitory concentrations of these inhibitors did not affect the growth of wild-type strain (Fig. 4d). We further corroborated our titan cell formation results with capsule analysis. As shown in the Fig. 4e and f, both CsA and FK506 treatment at sub-inhibitory concentrations resulted in reduction in titan cell associated capsule expansion (Supplementary Fig. 5b). Taken together our results strongly support the positive correlation between calcineurin pathway activity and titan cell formation in *C. neoformans*.

**Fig. 4.**
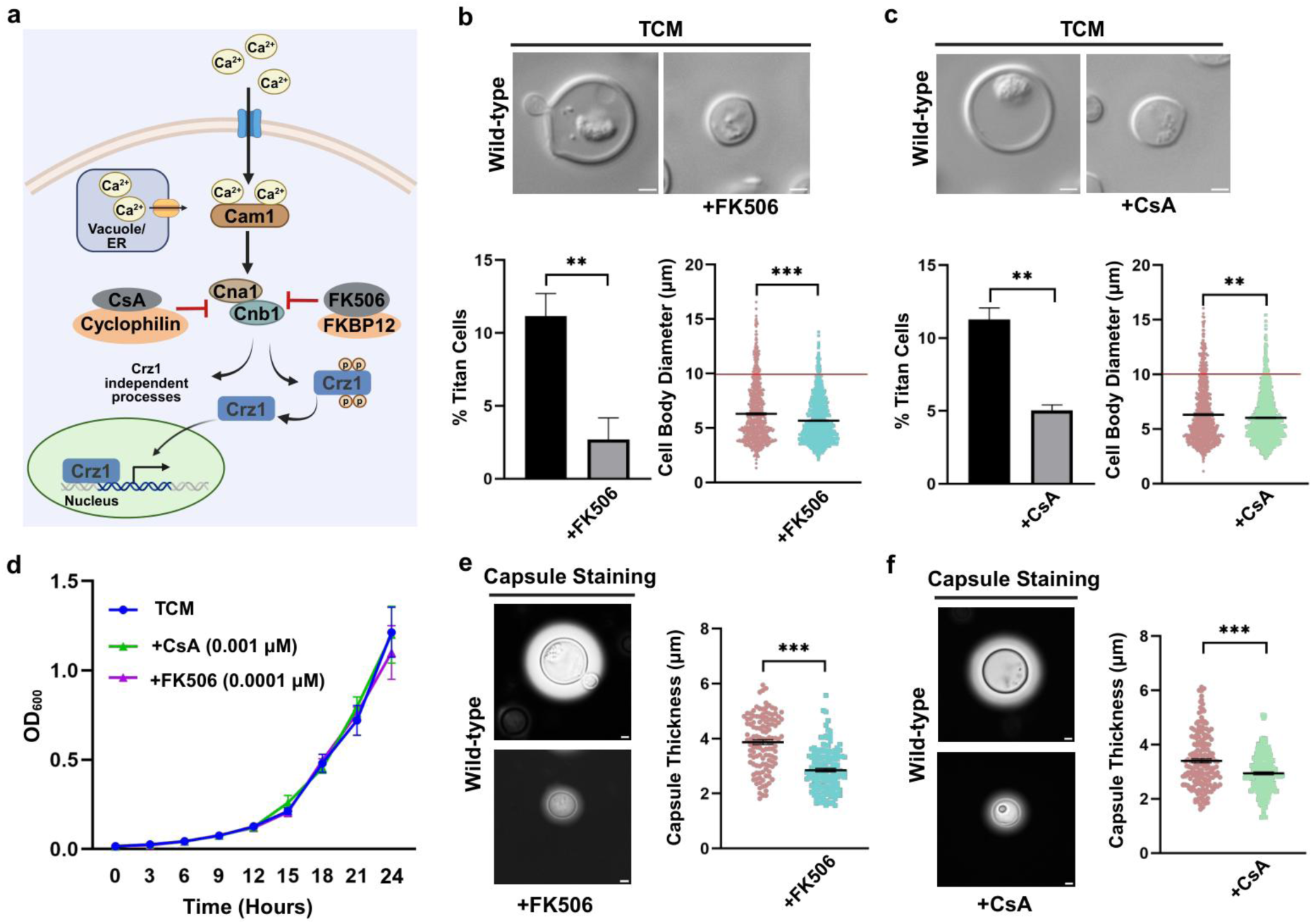
Inhibition of Calcineurin Impairs Titan Cell Formation in *C. neoformans.* a) Schematic overview of calcineurin pathway and its inhibition by inhibitors (Cyclosporine A (CsA) and FK506). b) Wild-type strain was cultured in Titan Cell Medium (TCM) with and without inhibitor (FK506 (0.0001 µM)) at 37 °C with 5% CO_2_ for 72 hours. Cells were fixed and imaged using Zeiss Apotome microscope. Percentage of titan cells was quantified as a ratio of cells with diameter ≥10 µm to total number of cells. Cell diameter was measured using Zen 2.3 software. Threshold for cell body diameter of titan cells is indicated by the red line. More than 400 cells were counted for each condition. Statistical analysis was performed using unpaired t-test, ***(P<0.001) and **(P<0.01). Error bars represent SEM. Scale bars represent 2 µm. c) Wild-type strain was cultured in TCM with and without inhibitor (CsA (0.001 µM)) at 37 °C with 5% CO_2_ for 72 hours. Cells were fixed and imaged using Zeiss Apotome microscope. Percentage of titan cells was quantified as a ratio of cells with a diameter ≥10 µm to total number of cells. Cell diameter was measured using Zen 2.3 software. Threshold for cell body diameter of titan cells is indicated by the red line. More than 400 cells were counted for each condition. Statistical analysis was performed using unpaired t- test, **(P<0.01). Error bars represent SEM. Scale bars represent 2 µm. d) Growth curve was performed to monitor overall growth of wild-type strain in TCM, + FK506 and +CsA conditions. Overnight grown culture of wild-type strain (H99) was diluted to OD_600_=0.01 in fresh TCM medium with and without FK506 (0.0001 µM) or CsA (0.001 µM) and allowed to grow at 37 °C for 24 hours. OD_600_ was recorded at 3-hour intervals. e) For capsule visualization, wild-type strain was cultured in TCM with and without FK506 at 37 °C with 5% CO_2_ for 72 hours, cells were stained with India ink and imaging was done using Zeiss Apotome microscope. Capsule thickness was measured using Zen 2.3 software. Approximately 150 cells were counted for each condition. Statistical analysis was performed using unpaired t-test, ***(P<0.001). Error bars represent SEM. Scale bars represent 2 µm. f) For capsule visualization, wild-type strain was cultured in TCM with and without CsA at 37 °C with 5% CO_2_ for 72 hours, cells were stained with India ink and imaging was done using Zeiss Apotome microscope. Capsule thickness was measured using Zen 2.3 software. Approximately 150 cells were counted for each condition. Statistical analysis was performed using unpaired t-test, ***(P<0.001). Error bars represent SEM. Scale bars represent 2 µm. This figure was created using BioRender. https://BioRender.com/be7g6rx.

### Glycolysis Sustains Calcium Homeostasis to Drive *C. neoformans* Titan Cell Formation

Our previous results suggest that glycolysis-mediated regulation of titan cell formation is, at least in part, dependent on calcineurin pathway activity. Since calcineurin functions as a calcium-dependent signaling pathway (Hemenway and Heitman 1999; Yadav and Heitman 2023), we hypothesized that glycolysis may influence calcineurin activity by altering intracellular calcium levels, which are essential for proper activation of the calcineurin pathway. To test this hypothesis, we measured intracellular calcium levels in wild-type and *hxk2*Δ mutant strains cultured under titan cell inducing conditions. Briefly, wild-type and *hxk2*Δ mutant strains were cultured under titan cell inducing conditions for 18 hours. To quantitatively assess intracellular calcium levels, cells were harvested following incubation and stained with the calcium-sensitive fluorescent dye Fura-2 AM. Intracellular calcium levels were determined using the ratiometric fluorescence measurement (340/380 nm), which provides a quantitative estimate of calcium concentration (Iida et al., 1990; Martínez et al., 2017). As shown in Fig. 5a, the *hxk2*Δ mutant exhibited significantly reduced intracellular calcium levels compared to the wild-type strain, indicating that glycolytic activity contributes to the maintenance of calcium homeostasis during titan cell induction.

**Fig. 5.**
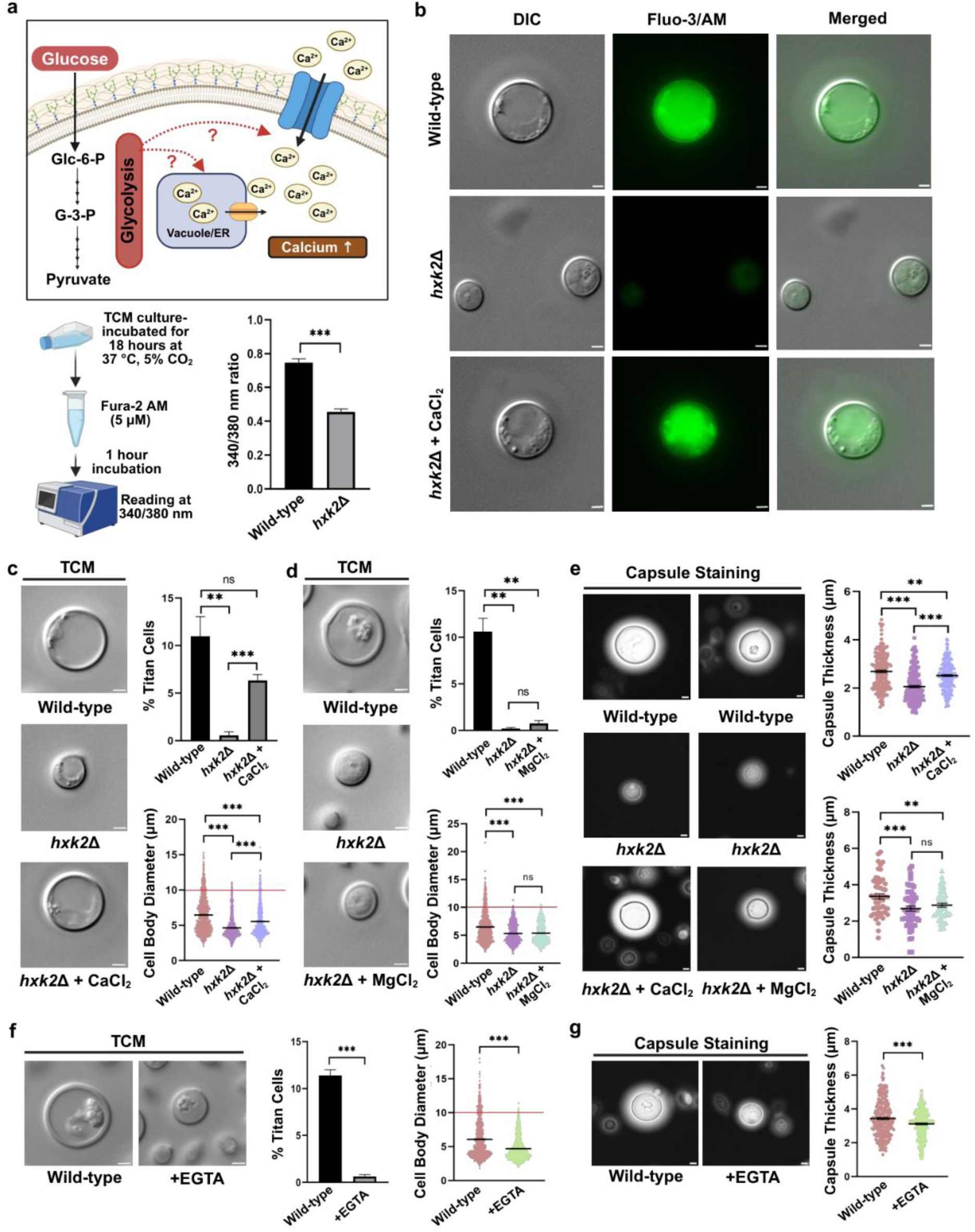
Glycolysis Sustains Calcium Homeostasis to Drive *C. neoformans* Titan Cell Formation. a) Quantitative measurement of intracellular calcium levels. Wild-type and *hxk2*Δ strain were cultured in titan cell inducing conditions for 24 hours. Cells were stained with calcium-sensitive dye, Fura-2 AM and fluorescence reading was taken as ratio 340/380 nm excitation and 510 nm emission. Statistical analysis was performed using unpaired t-test, ***(P<0.001). Error bars represent SEM. b) Qualitative measurement of intracellular calcium levels. Wild-type and *hxk2*Δ strain (with and without exogenous supplementation of CaCl_2_, at 2.5 mM) were cultured in titan cell inducing conditions for 24 hours. Cells were stained with calcium-sensitive dye, Fluo-3/AM (10 μM) and imaged in Zeiss Apotome microscope. Scale bars represent 2 µm. c) Wild-type and *hxk2*Δ strain were cultured in TCM with and without CaCl_2_ (2.5 mM) at 37 °C with 5% CO_2_ for 72 hours. Cells were fixed and imaged using Zeiss Apotome microscope. Percentage of titan cells was quantified as a ratio of cells with diameter ≥10 µm to total number of cells. Cell diameter was measured using Zen 2.3 software. Threshold for cell body diameter of titan cells is indicated by the red line. More than 400 cells were counted for each condition. Statistical analysis was performed using unpaired t-test, ***(P<0.001), **(P<0.01) and ns (non-significant). Error bars represent SEM. Scale bars represent 2 µm. d) Wild-type and *hxk2*Δ strain were cultured in TCM with and without MgCl_2_ (2.5 mM) at 37 °C with 5% CO_2_ for 72 hours. Cells were fixed and imaged using Zeiss Apotome microscope. Percentage of titan cells was quantified as a ratio of cells with a diameter ≥10 µm to total number of cells. Cell diameter was measured using Zen 2.3 software. Threshold for cell body diameter of titan cells is indicated by the red line. More than 400 cells were counted for each condition. Statistical analysis was performed using unpaired t-test, ***(P<0.001), **(P<0.01) and ns (non-significant). Error bars represent SEM. Scale bars represent 2 µm. e) For capsule visualization, wild- type and *hxk2*Δ strain were cultured in TCM with and without CaCl_2_ (2.5 mM) or MgCl_2_ (2.5 mM) at 37 °C with 5% CO_2_ for 72 hours, cells were stained with India ink and imaging was done using Zeiss Apotome microscope. Capsule thickness was measured using Zen 2.3 software. Approximately 150 cells were counted for each condition. Statistical analysis was performed using unpaired t-test, ***(P<0.001), **(P<0.01) and ns (non-significant). Error bars represent SEM. Scale bars represent 2 µm. f) Wild-type strain was cultured in TCM with and without EGTA (5 mM) at 37 °C with 5% CO_2_ for 72 hours. Cells were fixed and imaged using Zeiss Apotome microscope. Percentage of titan cells was quantified as a ratio of cells with a diameter ≥10 µm to total number of cells. Cell diameter was measured using Zen 2.3 software. Threshold for cell body diameter of titan cells is indicated by the red line. More than 400 cells were counted for each condition. Statistical analysis was performed using unpaired t-test, ***(P<0.001). Error bars represent SEM. Scale bars represent 2 µm. g) For capsule visualization wild-type strain was cultured in TCM with and without EGTA (5 mM) at 37 °C with 5% CO_2_ for 72 hours. Cells were fixed and imaged using Zeiss Apotome microscope. Cells were stained with India ink and imaging was done using Zeiss Apotome microscope. Capsule thickness was measured using Zen 2.3 software. Approximately 150 cells were counted for each condition. Statistical analysis was performed using unpaired t-test, ***(P<0.001). Error bars represent SEM. Scale bars represent 2 µm. This figure was created using BioRender. https://BioRender.com/jioidky.

In parallel, intracellular calcium levels were also qualitatively monitored using the calcium-sensitive fluorescent dye Fluo-3/AM in wild-type and *hxk2*Δ strains, with and without CaCl₂ supplementation (Fig. 5b) (Minta et al., 1989; Williams, 1990). Consistent with the Fura-2 AM results, the *hxk2*Δ mutant displayed reduced intracellular calcium levels relative to the wild-type. Importantly, exogenous CaCl₂ supplementation restored intracellular calcium levels in the *hxk2*Δ mutant, suggesting that the glycolytic mutant is impaired in maintaining calcium homeostasis efficiently under normal titan cell inducing conditions. Given this observation, we further assessed the effect of exogenous supplementation of calcium on overall titan cell formation in *hxk2*Δ in our in vitro titan cell formation assays. To confirm that any rescue observed was specifically due to calcium rather than a general effect of divalent cation supplementation, we also included MgCl₂ as a control. Wild-type and *hxk2*Δ mutant (in presence and absence of CaCl_2_/MgCl_2_) were cultured in titan cell inducing conditions. Cells were imaged using bright field microscopy, and percentage of titan cells was quantified as the ratio of titan cells (≥10 µm) to the total number of cells. As shown in Fig. 5c and d, exogenous CaCl₂ supplementation successfully rescued titan cell formation in the *hxk2*Δ mutant, whereas MgCl₂ had no such effect, confirming that the rescue is calcium-specific (Supplementary Fig. 6a). This finding was further corroborated by capsule analysis, wherein titan cell-associated capsule formation was similarly restored by CaCl₂ but not MgCl₂ supplementation in the *hxk2*Δ mutant (Fig. 5e; Supplementary Fig. 6b). Collectively, these results suggest that the reduction in intracellular calcium levels resulting from glycolytic disruption is a key contributor to the titan cell defect observed in the *hxk2*Δ mutant.

To further validate our hypothesis, we took a complementary approach and asked whether depleting extracellular calcium in the titan cell medium would affect the titan cell formation. In order to do this, we treated TCM media inoculated with wild-type *C. neoformans*, with EGTA, a well-established calcium chelator (Li et al. 2012; Liu et al. 2014) and titan cell formation was assessed. As shown in Fig. 5f and g (Supplementary Fig. 6a and b), EGTA treatment led to a significant reduction in titan cell formation and titan cell associated capsule expansion in the wild-type strain. This result provides independent support for the idea that calcium availability is a critical determinant of titan cell formation, and that the calcium deficit resulting from glycolytic disruption is likely a key driver of the impaired titanization observed in the *hxk2*Δ mutant.

### cAMP-PKA Signaling Acts as a Functional Bypass for Glycolytic Deficits by Restoring Calcium Homeostasis and Calcineurin Activity During Titanization

Our results clearly demonstrate that glycolysis-mediated calcium homeostasis and calcineurin activity play a critical role in titan cell formation. While the cAMP-PKA pathway is the most extensively studied signaling cascade regulating titan cell formation in *C. neoformans* (Alspaugh et al. 1997; Xue et al. 1998; Pukkila-Worley and Alspaugh 2004; Zaragoza et al. 2010), its relationship with calcium signaling remains nuanced. Although cAMP-PKA and calcineurin are generally considered independent, parallel signaling cascades, evidence across diverse biological systems suggests a positive correlation between cAMP-PKA and calcium signaling. This is supported by studies in *S. cerevisiae* showing that glucose addition induces transient increases in both cAMP levels and Ca²⁺ influx, suggesting a conserved link between cAMP signaling and calcium mobilization in fungi. (Eilam et al. 1990; Kamp and Hell 2000; Zaccolo et al. 2021). Therefore, we sought to determine whether activation of the cAMP-PKA pathway could restore intracellular calcium levels in our glycolysis-deficient mutant strain *hxk2*Δ. In order to test this hypothesis, we cultured *hxk2*Δ strain with or without exogenous supplementation of cAMP under titan cell inducing conditions for 18 hours and monitored intracellular calcium levels using calcium-sensitive dye Fluo-3/AM. Wild-type strain was used as a control. As shown in Fig. 6a, exogenous supplementation of cAMP, restored the intracellular calcium levels in *hxk2*Δ back to wild-type levels. Having observed that cAMP could rescue calcium levels in the glycolysis mutant, we sought to further validate our hypothesis and elucidate the underlying mechanistic details of cAMP-PKA-mediated calcineurin signaling during titan cell formation. To achieve this, we performed a comprehensive transcriptomic analysis comparing the wild-type and *hxk2*Δ mutant strains in the presence of exogenous cAMP, under titan cell inducing conditions. Additionally, we also compared our RNA-seq data set with the previously mentioned Chow et al. 2017 dataset to understand the effect of exogenous supplementation of cAMP on calcineurin pathway particularly (Chow et al. 2017). We observed that the expression of many of the calcineurin pathway genes which were previously found to be downregulated in our *hxk2*Δ mutant strain, were restored back to wild-type levels after exogenous supplementation of cAMP. This observation is illustrated using a volcano plots and Venn diagrams comparing all the known calcineurin pathway genes obtained from the dataset of Chow et al. 2017 with our RNA-seq dataset of wild-type and *hxk2*Δ, with and without exogenous supplementation of cAMP (Fig. 6b and c).

**Fig. 6.**
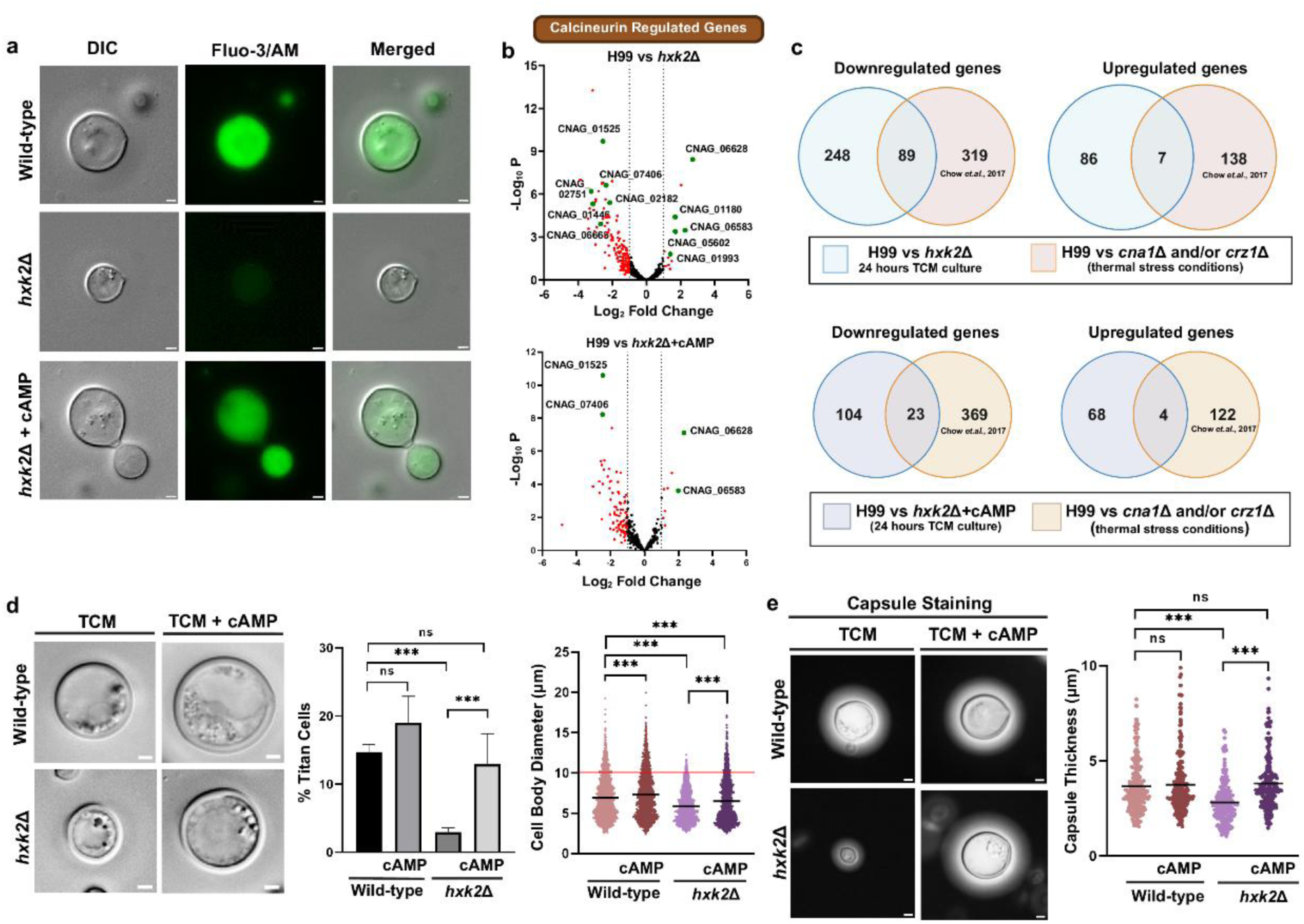
cAMP-PKA Signaling Acts as a Functional Bypass for Glycolytic Deficits by Restoring Calcium Homeostasis and Calcineurin Activity During Titanization. a) Qualitative measurement of intracellular calcium levels. Wild-type and *hxk2*Δ strain (with and without exogenous supplementation of cAMP, at 10 mM) were cultured in titan cell inducing conditions for 18 hours. Cells were stained with calcium-sensitive dye, Fluo-3/AM and imaged in Zeiss Apotome microscope. Scale bars represent 2 µm. b) Volcano plots represent calcineurin-regulated genes (which were identified by Chow et al. 2017) that are differentially expressed in *hxk*2Δ strain (with and without exogenous supplementation of cAMP (10 mM)) compared to wild-type strain grown under titan cell inducing conditions for 24 hours. The same wild-type replicates used for the wild-type vs *hxk2*Δ comparison (Fig. 3) were also used for the wild-type vs cAMP-supplemented *hxk2*Δ comparison. Genes which are upregulated (Log_2_ fold change ≥ 1) and downregulated (Log_2_ fold change ≤ -1) are highlighted in red. Some of the significantly upregulated and downregulated genes involved in the calcineurin pathway are highlighted in green. c) Venn diagram representing overlap between our RNA-seq dataset (wild-type and *hxk2*Δ (with and without exogenous supplementation of cAMP (10 mM)) under titan cell inducing condition) and dataset published by Chow et al. 2017, (wild-type vs calcineurin mutants under thermal stress condition). In order to highlight our observation that cAMP supplementation restores the expression of multiple calcineurin-regulated genes in *hxk2*Δ, the Venn diagram representing the overlap between our RNA-seq dataset (wild-type vs *hxk2*Δ under titan cell inducing condition) and dataset published by Chow et al. 2017 (wild-type vs calcineurin mutants under thermal stress condition) from Fig. 3d, has been reused in this panel. d) Wild-type and *hxk2*Δ strains were cultured in TCM in the presence and absence of cAMP (10 mM) at 37 °C with 5% CO_2_ for 72 hours. Cells were fixed and imaged using Zeiss Apotome microscope. Percentage of titan cells was quantified as a ratio of cells with a diameter ≥10 µm to total number of cells. Cell diameter was measured using Zen 2.3 software. Threshold for cell body diameter of titan cells is indicated by the red line. More than 400 cells were counted for each condition. Statistical analysis was performed using unpaired t-test, ***(P<0.001) and ns (non-significant). Error bars represent SEM. Scale bars represent 2 µm. e) For capsule visualization, wild-type and *hxk2*Δ strains were cultured in TCM in the presence and absence of cAMP (10 mM) at 37 °C with 5% CO_2_ for 72 hours. Cells were fixed and imaged using Zeiss Apotome microscope. Cells were stained with India ink and imaging was done using Zeiss Apotome microscope. Capsule thickness was measured using Zen 2.3 software. Approximately 150 cells were counted for each condition. Statistical analysis was performed using unpaired t-test, ***(P<0.001) and ns (non-significant). Error bars represent SEM. Scale bars represent 2 µm. This figure was created using BioRender. https://BioRender.com/lsjykca.

To validate our hypothesis further, we performed cAMP add-back assays wherein glycolysis knockout strains *hxk2*Δ with wild-type strain were cultured in TCM with and without the exogenous supplementation of cAMP (10 mM) and incubated under titan cell inducing conditions. Remarkably, exogenous supplementation of cAMP significantly rescued the titan cell formation defect observed in *hxk2*Δ (Fig. 6d; Supplementary Fig. 7a). The percentage of titan cells were significantly higher in the cAMP treated *hxk2*Δ strains compared to the untreated controls, reaching levels comparable to the wild-type strain. These results were further corroborated with our capsule analysis, wherein, addition of cAMP exogenously, restored the titan cell associated capsule formation in *hxk2*Δ mutant (Fig. 6e; Supplementary Fig. 7b).

## Discussion

Our findings provide compelling evidence for a previously uncharacterized regulatory axis linking central carbon metabolism, specifically glycolysis, and the calcineurin pathway in the regulation of titan cell formation in *C. neoformans*. We also demonstrate a possible connection between the cAMP-PKA pathway and calcium-calcineurin signaling through the maintenance of intracellular calcium levels, particularly in the context of titan cell formation. Despite the well-recognized importance of titan cells in the disease progression and pathogenesis of *C. neoformans* (Crabtree et al. 2012; Okagaki and Nielsen 2012), the mechanistic basis underlying this morphological transition remains poorly characterized. Titan cells exhibit dramatically altered cellular and morphological changes compared to the yeast morphotype, including an increase in cell size, polyploidy resulting from endoreduplication, and a thicker cellular envelope (cell wall and polysaccharide capsule) with altered composition (Zhou and Ballou 2018; García-Rodas et al. 2019). These pronounced morphological changes suggest that titan cells represent a metabolically active state of *C. neoformans*, triggered specifically under host-relevant conditions. While considerable efforts have been made to understand the metabolic adaptations of *C. neoformans* under host-relevant conditions (Lev et al. 2020), the relationship between cellular metabolism and the yeast-to-titan cell transition remains unexplored. Given that titan cell formation involves substantial biosynthetic requirements, including extensive remodelling of the cell wall and capsule, central carbon metabolism, and glycolysis in particular, emerges as a plausible metabolic node that is critical for this transition. Importantly, glycolysis plays a critical role in generating various precursors required for cell wall and capsule biosynthesis, and the role of glycolysis in regulating cell size and cell cycle progression in various organisms, including multiple fungal species (Cadart and Heald 2022; Diehl et al. 2024; Kierans and Taylor 2024). There is increasing evidence that suggests that glycolysis plays a regulatory role in orchestrating fungal morphogenesis in multiple fungal pathogens. Our recent work showed glycolysis orchestrates fungal morphogenesis in a sulfur-dependent manner in both *Saccharomyces cerevisiae* and the human fungal pathogen, *Candida albicans* (Shah et al. 2026). Similarly, several other studies have demonstrated the importance of glycolysis in the context of fungal morphogenesis although the mechanisms underlying these observations remain to be elucidated (Sabina and Brown 2009; Laurian et al. 2019). Interestingly, a recent study reported the successful generation of in vitro titan cells which are remarkably similar to the titan cells that are formed in vivo, during *C. neoformans* infections, demonstrated that in vitro titan cells have increased expression of several glycolytic genes (Trevijano-Contador et al. 2018). Given these observations, we hypothesized that glycolysis might play an important role in the regulation of titan cell formation in *C. neoformans*. Our results clearly demonstrated that the inhibition of glycolysis with sub-inhibitory concentrations of 2DG or NaCi significantly reduced titan cell formation, providing strong evidence for the metabolic dependency of this morphogenetic process. These findings were further supported by the results from titan cell assays using pertinent glycolysis deletion strains, wherein *hxk2*Δ (*hxk2*Δ refers to the *hxk2* deletion strain) showed a substantial reduction in titan cell formation compared to the wild-type.

Having established glycolysis as a key metabolic driver of titan cell formation, we next sought to identify the downstream signaling mechanisms through which glycolytic flux might orchestrate this morphogenetic transition. Transcriptomic comparison between the wild-type and *hxk2*Δ strains under titan cell inducing conditions provided a critical and unexpected insight. A significant proportion of genes downregulated in the glycolysis-deficient mutant overlapped with genes identified within the calcineurin-responsive dataset reported by Chow et al. (2017), suggesting that calcineurin signaling may function as a downstream mediator of glycolytic regulation during titanization. In *C. neoformans*, the calcineurin pathway represents one of the most extensively characterized signaling cascades governing fungal virulence, mediating adaptive responses to host-relevant stresses such as physiological temperature, elevated CO₂, and pH fluctuations, alongside other well-characterized pathways including cAMP-PKA and Cell Wall Integrity signaling (Pukkila-Worley and Alspaugh 2004; Liu et al. 2014; Chow et al. 2017; Mazzi et al. 2018; Squizani et al. 2020; Squizani et al. 2021; Yadav and Heitman 2023). However, despite its well-established role in stress adaptation and virulence, a direct involvement of calcineurin signaling in titan cell formation had not previously been investigated, making our observation particularly significant. This was functionally corroborated by pharmacological inhibition of calcineurin using CsA and FK506, both of which significantly attenuated titan cell formation at sub-inhibitory concentrations without compromising cell growth (Chow et al. 2017; Kalem et al. 2021; Yadav and Heitman 2023), collectively establishing calcineurin as a positive regulator of titan cell formation. Mechanistically, our data suggest that glycolysis sustains calcineurin pathway activity by maintaining intracellular calcium homeostasis. Supporting this, *hxk2*Δ mutants exhibited markedly reduced intracellular calcium levels, and exogenous CaCl₂ supplementation was sufficient to rescue both intracellular calcium levels and titan cell formation, positioning calcium availability as a pivotal node connecting central carbon metabolism to morphogenetic signaling. The specificity of this rescue, observed with CaCl₂ but not with the divalent cation control MgCl₂, reinforces the conclusion that the glycolysis-calcineurin crosstalk is calcium-dependent, rather than a reflection of a general ionic effect. Further substantiating this mechanistic model, chelation of extracellular calcium using EGTA (Liu et al. 2014) significantly impaired titan cell formation in the wild-type strain, underscoring the indispensable role of calcium availability in this morphogenetic programme.

This prompted us to consider how glycolytic activity might interface with the calcium regulatory machinery in fungi, several components of which are unique to the fungal kingdom. In fungi, intracellular calcium homeostasis is maintained through the coordinated activity of a repertoire of transporters, channels, and pumps distributed across the plasma membrane and intracellular organelles, including the vacuole and endoplasmic reticulum (ER) (Hemenway and Heitman 1999; Yadav and Heitman 2023). Under resting conditions, cytosolic calcium is actively maintained at nanomolar concentrations through sequestration into intracellular stores by SERCA-type pumps and extrusion via P-type Ca²⁺-ATPases such as Pmr1 (Liu et al. 2015; Yadav and Heitman 2023; Lukumay et al. 2025). At the plasma membrane, calcium influx is primarily mediated by the Mid1-Cch1 complex, a fungal-specific high-affinity calcium channel that plays a critical role in calcium uptake in response to a wide range of stress stimuli, including membrane stretch, azole antifungals, and alkaline pH (Liu et al. 2006; Hong et al. 2013; Vu et al. 2015). Within the cell, the vacuole serves as the predominant calcium reservoir in fungi, with calcium sequestration being mediated by the Pmc1 P-type Ca²⁺-ATPase and the Vcx1 vacuolar H⁺/Ca²⁺ exchanger (Cunningham and Fink 1994; Kmetzsch et al. 2010; Cunningham 2011; Liu et al. 2015; Yadav and Heitman 2023). Calcium release from the vacuole is primarily governed by Yvc1, a fungal TRP-like channel that responds to membrane stretch, osmotic stress, and elevated cytosolic ROS, and whose proper assembly and vacuolar localisation depends on V-ATPase-driven vacuolar acidification (Marsh 2001; Peng et al. 2020). Upon stress stimulation, the concerted action of these fungal calcium uptake and release systems drives a transient rise in cytosolic calcium, which is sensed by calmodulin. Calmodulin, upon calcium binding, undergoes a conformational change that directly activates calcineurin, initiating downstream transcriptional responses through dephosphorylation of effectors such as Crz1 and Lhp1 (Park et al. 2019; Squizani et al. 2021; Yadav and Heitman 2023). Critically, all of the calcium transporters, exchangers, and ATPase pumps described above are dependent actively or passively on ATP availability. This creates a direct functional dependency between glycolytic flux and calcium homeostasis: given that glycolysis represents a primary source of cytosolic ATP, any perturbation in glycolytic activity would be expected to compromise the energetics of calcium transport, thereby disrupting the regulated calcium transients required for calcineurin activation. Consistent with this reasoning, glucose-starved cells have previously been shown to exhibit calcium uptake upon glucose re-addition via glucose-induced calcium (GIC) systems (Eilam et al. 1990; Groppi et al. 2011), highlighting the tight coupling between glycolytic activity and calcium mobilisation. In this aspect, the markedly reduced intracellular calcium levels observed in *hxk2*Δ mutant under titan cell inducing conditions could be attributed to impaired ATP-dependent calcium uptake or mobilisation resulting from reduced glycolytic output.

Beyond bioenergetic support, glycolysis may further regulate calcium dynamics through direct metabolite-mediated mechanisms. Glycolytic intermediates such as glucose-6-phosphate have been reported to modulate intracellular calcium homeostasis in yeast (Aiello et al. 2002), while the proper glucose-dependent assembly and activity of the V-ATPase, which drives vacuolar acidification essential for Yvc1-mediated calcium release, has been shown to depend on direct physical interactions with glycolytic enzymes including aldolase and phosphofructokinase-1 (Aiello et al. 2002; Lu et al. 2004; Chan and Parra 2014; Peng et al. 2020). Also, DAG, a lipid mediator biosynthetically linked to the glycolytic intermediate DHAP via the glycerophospholipid synthesis pathway, modulates calcium signaling through direct activation of TRP-type channels and PKC-mediated signaling, while its co-production with IP3 via phospholipase C-mediated PIP2 hydrolysis couples DAG signaling to IP3-dependent calcium release from intracellular stores (Hofmann et al. 1999; Berridge et al. 2000). The dependency of Yvc1 assembly and vacuolar localisation on V-ATPase activity further links glycolytic flux directly to fungal-specific vacuolar calcium release machinery (Peng et al. 2020). Extending this further, our transcriptomic analysis identified a gene encoding glutathione-S-transferase as significantly downregulated in the *hxk2*Δ mutant (Log_2_ fold change -2.554). This enzyme plays a particularly important role under elevated ROS conditions, where it catalyses the glutathionylation of Yvc1, activating calcium release into the cytosol and engaging calcineurin signaling (Chandel et al. 2016). Notably, titanization is known to occur under conditions of elevated endogenous ROS, which may itself serve as a trigger for calcineurin activation through Yvc1-mediated calcium mobilisation (Chandel et al. 2016; García-Barbazán et al. 2024). Downregulation of glutathione-S-transferase in the glycolysis-deficient mutant would therefore be expected to impair this ROS-responsive Yvc1 activation, further contributing to the reduced intracellular calcium levels and attenuated calcineurin signaling observed in this strain. Taken together, these observations indicate that glycolysis sustains the calcium transients necessary for calcineurin activation through multiple convergent mechanisms, encompassing bioenergetic support of fungal calcium ATPases and channels, metabolite-level modulation of calcium transport machinery, and ROS-responsive regulation of vacuolar calcium release via Yvc1. This positions glycolytic flux as a central integrator that couples the metabolic state of the cell to morphogenetic signaling during titan cell development in *C. neoformans*. Further studies are warranted to delineate the exact mechanisms underlying glycolysis-mediated regulation of calcineurin activation in *C. neoformans*.

In mammalian systems, cAMP-PKA signaling has been shown to directly modulate the activity of plasma membrane calcium channels and intracellular calcium mobilisation machinery, positioning it as a positive upstream regulator of intracellular calcium availability (Kamp and Hell 2000; Zaccolo et al. 2021). This is supported by a study that demonstrated that a transient rise in cellular cAMP levels has been observed in *S. cerevisiae* following glucose addition, coinciding with a rapid transient increase in Ca²⁺ influx across the plasma membrane, suggesting that the coupling between cAMP signaling and calcium mobilisation is a conserved feature of fungal biology rather than a mammalian-specific phenomenon (Eilam et al. 1990). Our findings suggest that an analogous functional relationship operates in *C. neoformans* in the context of titanization. Exogenous cAMP supplementation in the glycolysis-deficient *hxk2*Δ mutant restored intracellular calcium levels and significantly restored calcineurin pathway gene expression under titan cell inducing conditions, suggesting that the cAMP-PKA pathway, the most extensively characterized regulator of titan cell development in *C. neoformans*, may function not merely as an independent morphogenetic switch, but also as a modulator of calcium homeostasis, thereby indirectly impinging on calcineurin pathway activity. One plausible mechanistic basis for this is that the PKA-mediated phosphorylation and activation of fungal calcium channels and transporters, including components of the Mid1-Cch1 complex and vacuolar calcium release machinery such as Yvc1, which could promote calcium influx and mobilisation could be contributing to the activation of calcineurin signaling under titan cell inducing conditions. Taken together, these observations point towards a functional intersection between the cAMP-PKA and calcium-calcineurin pathways, mediated at least in part through the regulation of intracellular calcium availability, and place glycolytic flux as a central coordinator of this signaling crosstalk during titan cell development in *C. neoformans*

In *C. neoformans*, the calcineurin signaling pathway plays a pivotal role in enabling fungal survival within the host environment, primarily by regulating cell wall integrity in response to a broad range of stress stimuli encountered during infection (Cunningham 2011; Squizani et al. 2021; Yadav and Heitman 2023). This pathway operates through both Crz1-dependent transcriptional mechanisms, driving expression of cell wall remodelling enzymes such as the chitin synthases, calcium transporters, and through Crz1-independent post-transcriptional mechanisms involving RNA-binding proteins such as Pbp1, Puf4, and Lhp1, underscoring the complexity and adaptability of calcineurin signaling in this pathogen (Chow et al. 2017; Yadav and Heitman 2023). Titan cell formation is a striking morphogenetic transition induced under host-relevant conditions, including elevated temperature, elevated CO₂, nutrient limitation, and serum exposure, precisely the conditions under which calcineurin activity is known to be elevated (Zaragoza et al. 2010; Zhou and Ballou 2018; García-Rodas et al. 2019; Yadav and Heitman 2023; Mahmood et al. 2024). Titanization is accompanied by profound structural remodelling, including dramatic enlargement of the cell body and vacuole, thickening and reorganisation of the cell wall, and extensive capsule expansion (Zaragoza et al. 2010; Zhou and Ballou 2018; García-Rodas et al. 2019). Given that calcineurin is one of the central pathways activated under these host-mimicking conditions, and that it directly governs the transcriptional programmes responsible for cell wall biosynthesis and stress adaptation (Squizani et al. 2020; Yadav and Heitman 2023; Mahmood et al. 2024), it is plausible that calcineurin plays a regulatory role in orchestrating the cellular remodelling events associated with titan cell development. Consistent with this, our transcriptomic data revealed significant downregulation of several calcineurin-regulated genes involved in thermal stress response, carbohydrate metabolism, and cell wall remodelling, including a glucosidase (CNAG_06835), in the *hxk2*Δ mutant (Table 1). Given that titanization involves dramatic cell size expansion accompanied by extensive cell wall remodelling, the downregulation of genes governing carbohydrate metabolism and cell wall integrity in *hxk2*Δ likely underlies its titan cell defect, further implicating active calcineurin signaling as a critical mediator of titan cell formation. Further supporting a role for calcineurin in titanization, our transcriptomic analysis also identified significant downregulation of *RBD*1 (Log_2_ fold change -1.7225) in the *hxk2*Δ mutant, a gene previously shown to be required for efficient titan cell formation (Shu et al. 2026), providing an additional mechanistic link between impaired calcineurin pathway activity and the reduction in titanization observed in this strain. However, the precise mechanism by which calcineurin influences titanization remains to be fully elucidated. A key unresolved question is whether calcineurin exerts its effect on titan cell formation through its canonical downstream effector Crz1, or through one of its Crz1-independent branches. These two possibilities have distinct implications: if the effect is Crz1-dependent, it would implicate a transcriptional programme driven by CDRE-containing target gene promoters; if it is Crz1-independent, it would instead point toward post-transcriptional regulation or other non-transcriptional calcineurin substrates as the relevant effectors of morphogenesis (Chow et al. 2017). A direct and informative approach to address this question would be to examine the subcellular localisation of Crz1 under titan cell inducing conditions. Since calcineurin-dependent activation of Crz1 is manifested by its dephosphorylation and subsequent nuclear translocation (Gupta et al. 2022), monitoring Crz1 localisation in titan cell inducing media would reveal whether calcineurin is indeed signaling through Crz1 during this morphogenetic transition. Nuclear accumulation of Crz1 under these conditions would strongly suggest a Crz1-dependent mechanism, whereas cytoplasmic retention despite calcineurin activation would point toward Crz1-independent effectors as the primary mediators. Additionally, testing a *crz1* knockout strain (*crz1*Δ) for its ability to form titan cells under in vitro titanization assays would provide valuable information about the contribution of Crz1 towards this morphological transition. Complementarily, to determine whether calcineurin influences titanization through its Crz1- independent arm, which primarily involves regulation of P-bodies and RNA metabolism, examining the localisation of the calcineurin catalytic subunit Cna1 would be informative. Previous studies have demonstrated that calcineurin exerts Crz1-independent regulatory roles by relocalising to ER sites or P-bodies (Kozubowski et al. 2011; Chow et al. 2017), and determining whether similar redistribution occurs under titan cell inducing conditions would shed light on the non-transcriptional mechanisms through which calcineurin may govern this morphogenetic transition. Such experiments would provide a critical framework for delineating the downstream architecture of calcineurin signaling during titan cell development in *C. neoformans*, and may ultimately reveal novel regulatory nodes linking stress signaling to fungal morphogenesis within the host.

In summary, our study uncovers a previously uncharacterized regulatory axis linking central carbon metabolism to morphogenetic signaling in *C. neoformans*, wherein glycolytic flux serves as a critical upstream determinant of titan cell formation (Fig. 7). Mechanistically, we demonstrate that glycolysis sustains intracellular calcium homeostasis possibly through the modulation of ATP-dependent calcium transport machinery, thereby maintaining the calcium transients necessary for calcineurin pathway activation and downstream morphogenetic gene expression. Furthermore, our findings reveal that the cAMP-PKA pathway, the canonical regulator of titanization, intersects with this glycolysis-calcium-calcineurin axis by modulating intracellular calcium availability, suggesting that these signaling cascades operate in a functionally integrated rather than strictly independent manner during titan cell formation. Collectively, our work positions glycolytic flux as a metabolic checkpoint that coordinates energy availability, calcium homeostasis, and stress-responsive signaling to drive titan cell morphogenesis, and opens new avenues for understanding how the metabolic state of *C. neoformans* shapes its virulence potential within the host.

**Fig. 7.**
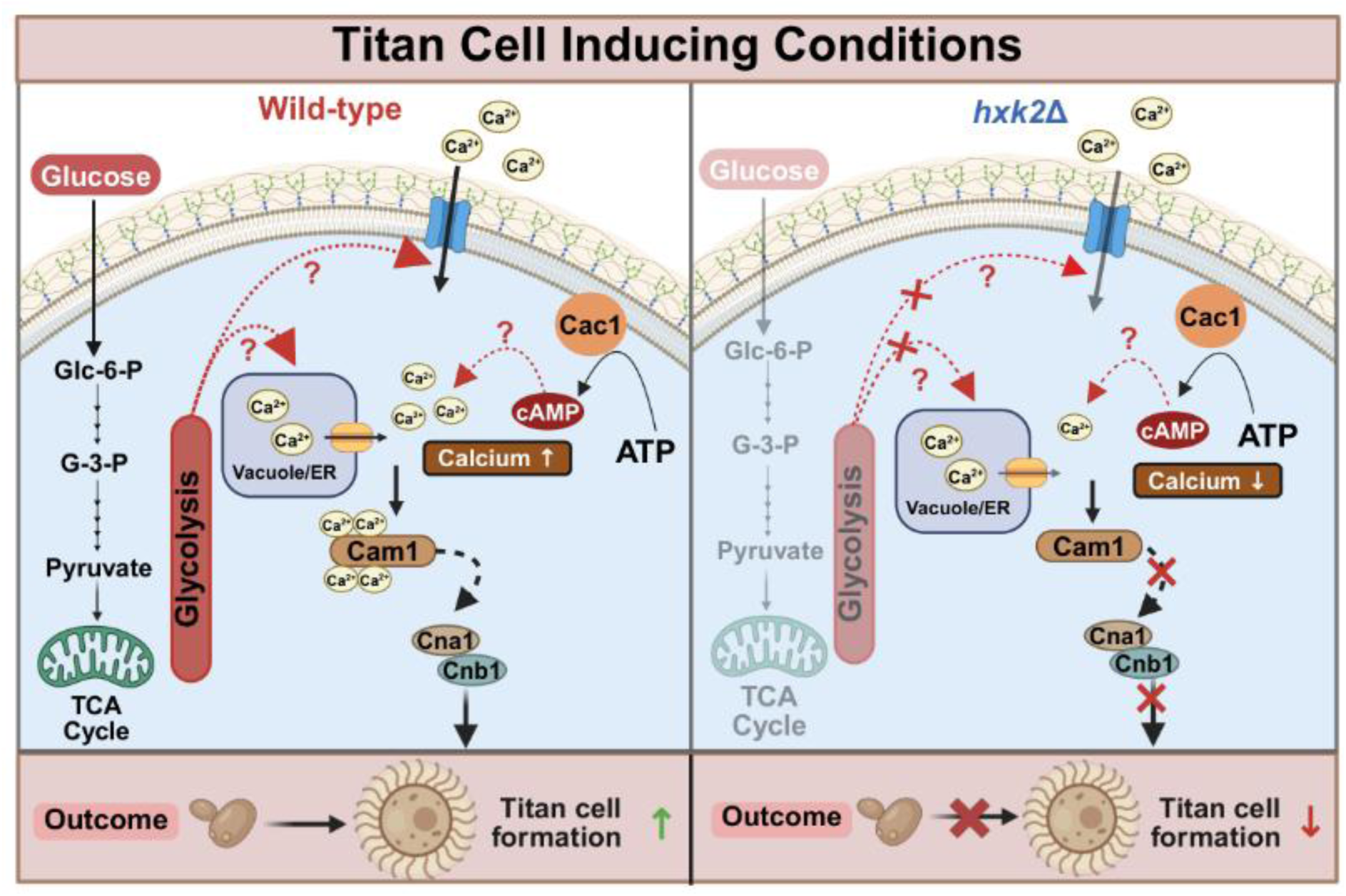
Glycolytic Regulation of Calcium Homeostasis and Calcineurin Signaling During Titan Cell Formation in *C. neoformans*. Under titan cell inducing conditions, active glycolysis maintains intracellular calcium homeostasis, promoting activation of the calcineurin pathway, which leads to titan cell formation. In the *hxk2*Δ mutant, impaired glycolysis disrupts calcium homeostasis, reduces calcineurin activation, and compromises titan cell formation. This figure was created using BioRender. https://BioRender.com/e17d3po.

## Materials and Methods

### Strains and Media

All strains used in this study are listed in Table 2. All experiments were performed using *Cryptococcus neoformans* var. *grubii*, serotype A, H99 wild-type, and its derivatives. All strains used in this study were generously provided by Dr. Joeseph Heitman, Duke University, USA. Yeast Peptone (YP) (1% (w/v) yeast extract and 2% (w/v) peptone) medium supplemented with glycerol was used for cryo-preservation of strains at -80 °C. All the strains were recovered at 30 °C on YPD agar plates (1% (w/v) yeast extract, 2% (w/v) peptone, 2% (w/v) glucose, and 2% (w/v) agar).

**Table 2:**
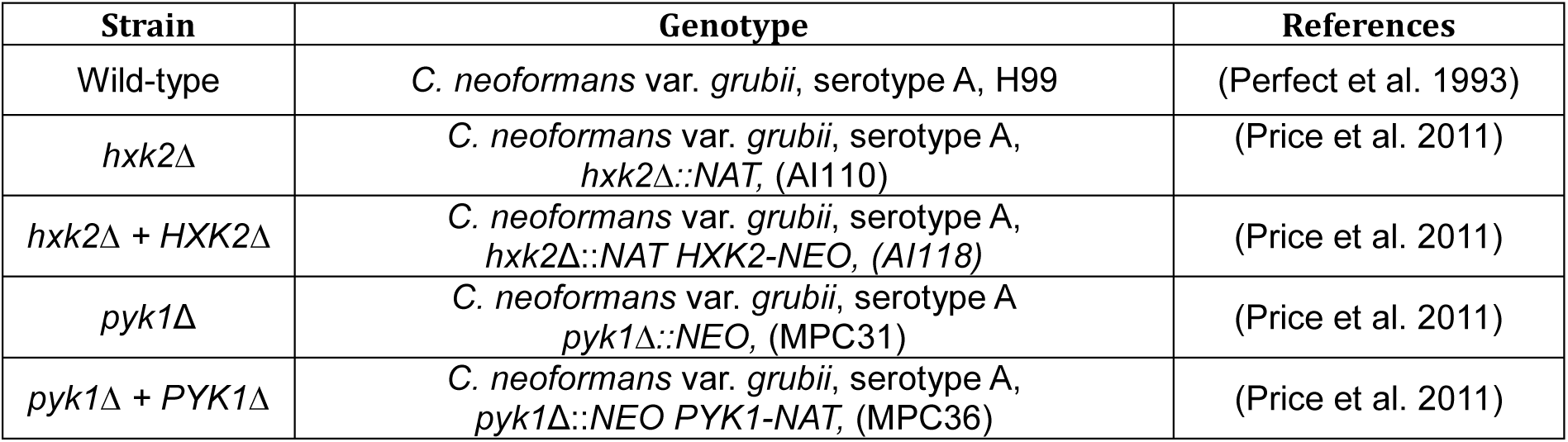
*C. neoformans* Strains Used in This Study.

YPD broth containing 1% (w/v) yeast extract, 2% (w/v) peptone, and 2% (w/v) glucose was used for the overnight growth of the cultures. Overnight cultures of various strains were grown in YPD broth and incubated at 30 °C with continuous shaking at 200 Rotations Per Minute (RPM). Titan Cell Medium (TCM) containing 5% (w/v) SDA (Sabouraud) (cat. #238230), 5% (v/v) FBS (Fetal Bovine Serum) (cat. #16000044), 15 µM Sodium Azide (cat. #S2002), and 50 mM MOPS (cat. #M3183) was used for all in vitro titan cell assays.

### In Vitro Titan Cell Assays

Overnight cultures of *C. neoformans* strains (listed in Table 2) were inoculated into Titan Cell Inducing Medium (TCM) at a standardized cell density of 10^6^ CFU per ml for 72 hours at 37 °C under 5% CO_2_.

#### Glycolysis inhibitor assays

To assess the effect of glycolysis perturbation on titan cell formation, cultures were grown in TCM supplemented with 2-Deoxy-D-Glucose (2DG; 0.025% w/v) (cat. #D8375) or sodium citrate (NaCi; 0.5% w/v) (cat. #C8532) for 72 hours under the standard conditions described above.

#### Calcineurin inhibitor assays

To determine the contribution of calcineurin signaling to titan cell induction, cultures were grown in TCM supplemented with FK506 at concentrations of 0.001 µM or 0.0001 µM, (cat. #F4679) or with cyclosporin A (CsA) at concentrations of 0.01 µM or 0.001 µM (cat. #C1832), for 72 hours under standard conditions.

#### Divalent cation supplementation assays

To examine the role of divalent cations in titan cell formation, wild-type and *hxk2Δ* strains were cultured in TCM in the presence or absence of CaCl₂ (2.5 mM) (cat. #0344317) or MgCl₂ (cat. #91417) (2.5 mM) for 72 hours under standard conditions.

#### EGTA chelation assays

To assess the effect of extracellular calcium depletion on titan cell induction, the wild-type strain was cultured in TCM supplemented with the calcium chelator EGTA (5 mM) (cat. #MB130) for 72 hours under standard conditions.

#### cAMP add-back assays

To determine whether exogenous cAMP can rescue titan cell defects caused by glycolysis perturbation, wild-type and *hxk2Δ* strains were cultured in TCM supplemented with cAMP (10 mM) (cat. #A9501) for 72 hours under standard conditions.

#### Imaging, quantification, and statistical analysis

Following 72 hours of incubation under the respective conditions, cells were fixed and imaged using a Zeiss Apotome fluorescence microscope. Cell diameter was measured using ZEN 2.3 software. Titan cell quantification was performed on a minimum of 400 cells per sample per condition, and titan cell percentage was defined as the proportion of cells with a cell body diameter of ≥10 µm relative to the total number of cells counted. Statistical analysis was performed using unpaired t-test (GraphPad Prism (v 9.0)). Error bars represent the Standard Error of the Mean (SEM). All statistical comparisons were based on a minimum of three independent biological replicates.

### Capsule Visualization and Analysis

For visualization and analysis of capsule thickness, overnight grown cultures of different *C. neoformans* strains (listed in Table 2) were inoculated in TCM at cell density of 10^6^ CFU per ml for 72 hours at 37 °C under 5% CO_2_. Following 72 hours of incubation, cells were fixed and stained using India ink (cat. #S025). Samples were imaged using Zeiss Apotome microscope. Capsule thickness was measured using Zen 2.3 software (150 cells each sample). Statistical analysis was performed using unpaired t-test (GraphPad Prism (v 9.0)). Error bars represent the Standard Error of the Mean (SEM). All statistical comparisons were based on a minimum of three independent biological replicates.

### Intracellular Calcium Level Measurement

Overnight cultures of the wild-type and *hxk2*Δ strains of *C. neoformans* were inoculated in TCM at cell density of 10^6^ CFU per ml and incubated for 18 hours at 37 °C under 5% CO₂ to induce titan cell formation. For experiments involving calcium supplementation, 2.5 mM CaCl₂ was added to the culture medium during incubation. Following incubation, cells were harvested by centrifugation, washed twice with 1X PBS, and resuspended in 1 ml of 1X PBS. For quantitative estimation of intracellular calcium levels, cells were treated with the calcium-sensing fluorescent dye Fura-2 AM (cat. #F1221) at a final concentration of 5 µM and incubated at 37 °C in the dark. After washing to remove excess dye, fluorescence was measured at 340 nm and 380 nm excitation with 510 nm emission, and intracellular calcium levels were calculated by dividing the 340/380 fluorescence intensity ratio. For qualitative assessment of intracellular calcium levels, cells were stained with the calcium-sensing fluorescent dye Fluo-3/AM (cat. #343242) at a final concentration of 10 µM under identical incubation conditions. Fluorescence images were taken using Zeiss Apotome microscope. Statistical analysis was performed using unpaired t-test (GraphPad Prism (v 9.0)). Error bars represent the Standard Error of the Mean (SEM). All statistical comparisons were based on a minimum of three independent biological replicates.

### Growth Curve Experiments

Overnight cultures of wild-type and knockout strains of *C. neoformans* (listed in Table 2) were inoculated in TCM at OD_600_ of 0.01 and incubated at 37 °C under continuous shaking conditions at 200 RPM for 24 hours. OD_600_ was recorded at 3-hour intervals. For growth curve analysis in the presence of glycolysis inhibitors, overnight cultures were inoculated in TCM in the presence and absence of 0.025% (w/v) of 2-Deoxy-D-Glucose (2DG) or 0.5% (w/v) of Sodium citrate (NaCi) and incubated at 37°C under continuous shaking conditions at 200 RPM for 24 hours. OD_600_ was recorded at 3-hour intervals. Experiments were performed in three independent replicates. For growth curve analysis in the presence of calcineurin inhibitors, overnight cultures were inoculated in TCM in the presence and absence of 0.001 µM and 0.0001 µM of FK506 and 0.01 µM and 0.001 µM of CsA and incubated at 37 °C under continuous shaking conditions at 200 RPM for 24 hours. OD_600_ was recorded at 3-hour intervals. Experiments were performed in three independent replicates.

### RNA Preparation for RNA-seq and Transcriptomic Data Analysis

Wild-type, *hxk2*Δ (with and without exogenous supplementation of cAMP) strains of *C. neoformans* were cultured in TCM at cell density of 10^6^ CFU per ml for 24 hours at 37 °C under 5% CO_2_. Total RNA was isolated from cells harvested after 24 hours of incubation using the hot acid-phenol method. For cell disruption, zirconium beads were used with a bead-beating protocol consisting of 10-second pulses interspersed with 1-minute rests on ice to prevent thermal degradation.

### RNA-sequencing analysis

Raw sequencing reads were obtained in FASTQ format. The reference genome and annotation files for the *C. neoformans* var. *grubii* H99 were downloaded from the NCBI site (Sample ID: SAMN03081441). Quality assessment of the raw reads was performed using FastQC (v 0.12.1). Adapter sequences were trimmed using Cutadapt (v 5.2). The reference genome was indexed with Hisat2 (v 2.2.1), and Samtools (v 1.20) was used to filter out multi-mapped reads and convert SAM files to BAM format. Read counting for uniquely mapped reads was done with FeatureCounts, using gene annotation data from the reference *C. neoformans* var. *grubii* H99 genome. Normalization and differential gene expression analysis were performed using R (v 4.3.2). Genes with a log_2_ fold change of ≥ 1 were considered to be upregulated, indicating at least a doubling in expression under the given condition, while those with a log_2_ fold change of ≤ -1 were considered to be downregulated.

### Reverse Transcriptase Quantitative Polymerase Chain Reaction (RT-qPCR) Assay

Wild-type and *hxk2*Δ strains of *C. neoformans* were cultured in TCM at cell density of 10^6^ CFU per ml for 24 hours at 37 °C under 5% CO_2_. Total RNA was extracted following 72 hours of incubation using the hot-phenol method. cDNA biosynthesis was done using PrimeScript cDNA biosynthesis kit (cat. #6110A). Primers used for RT-qPCR were designed using the PrimerQuest tool from Integrated DNA Technologies (IDT), and the primers used in these experiments are listed in Table 3. RT-qPCR was performed using iTaq Universal SYBR Green Master Mix (cat. #1725121). Actin was used as housekeeping gene for normalization. The results were analysed using the 2*^-ΔΔCT^*method and statistical analysis was performed using unpaired t-test (GraphPad Prism (v 9.0)). Error bars represent the Standard Error of the Mean (SEM). All statistical comparisons were based on a minimum of three independent biological replicates.

**Table 3:**
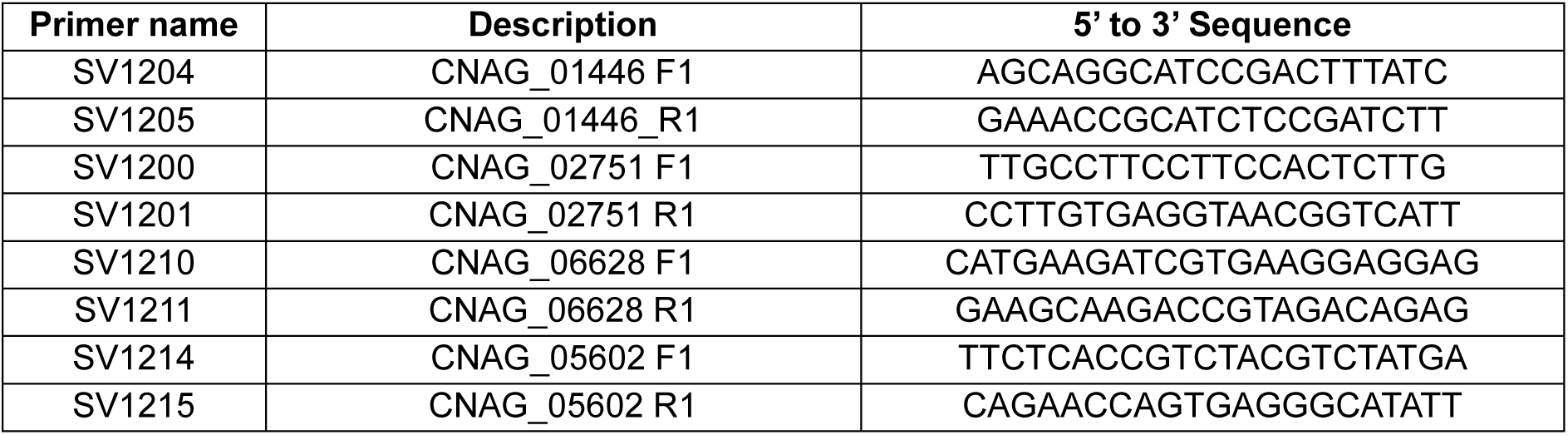
Primers Used in This Study.

### Illustrations

Figure illustrations were created using BioRender (https://app.biorender.com).

## Data Availability

The authors affirm that all data necessary for confirming the conclusions of the article are present within the article, figures, and Supplementary materials. RNA-seq data have been deposited in the NCBI SRA database (BioProject accession number PRJNA1501675) and in the NCBI GEO database (Accession number GSE341520).

## Acknowledgements

We extend our immense gratitude to Dr. Joseph Heitman and Mrs. Anna Floyd-Averette for generously providing the strains used in this study. We would like to thank Dr. Sunil Laxman (BRIC-inStem) and Dr. Vikas Yadav (Duke University) for critical reading of our manuscript. We gratefully acknowledge the invaluable support and resources offered by CSIR-Centre for Cellular and Molecular Biology (CCMB) central facilities including Advanced Microscopy and Imaging facility (AMIF). PSP thanks Council of Scientific and Industrial Research (CSIR), India for her research fellowship.

---

**Supplementary Figure 1.**
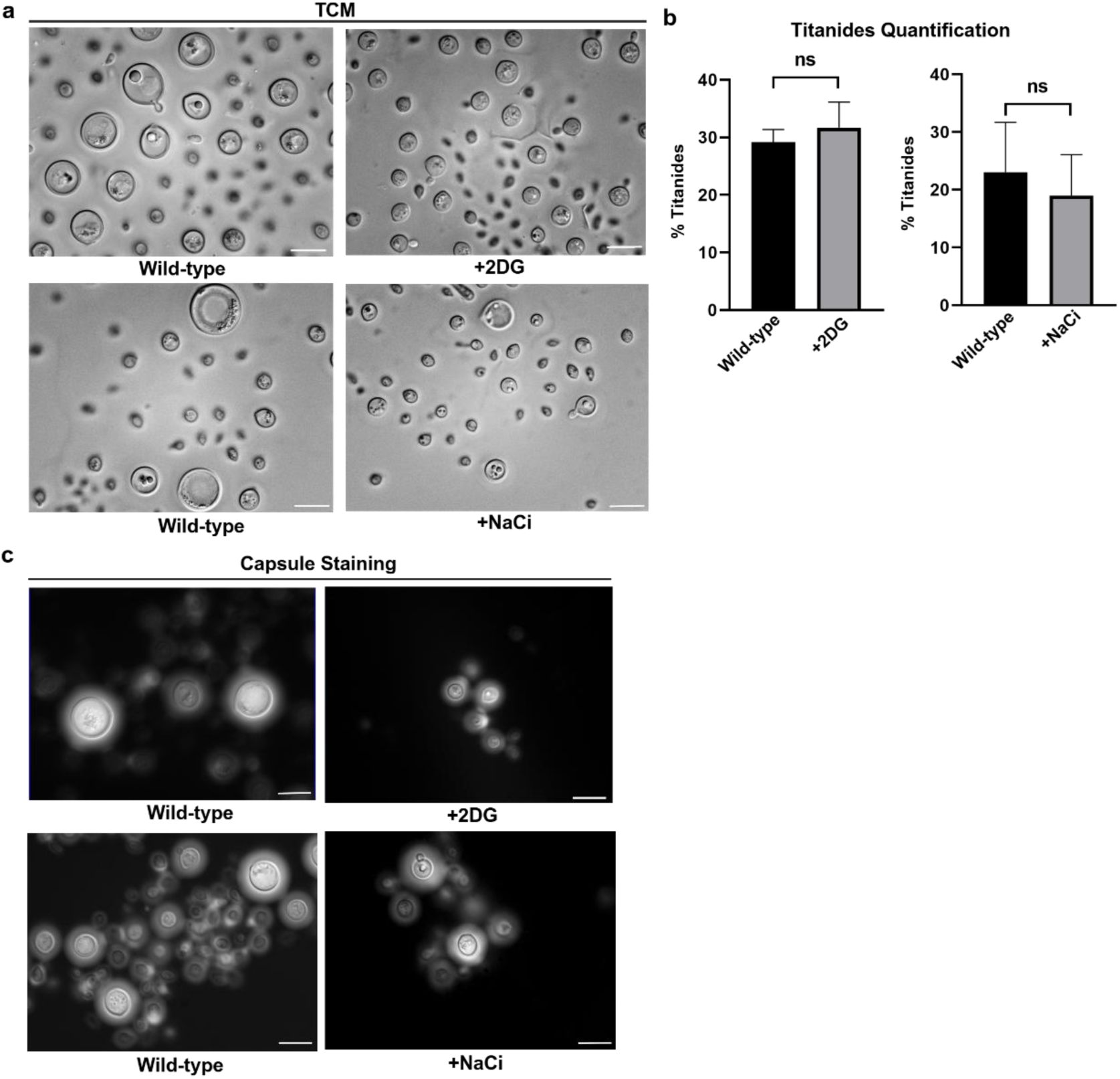
Wider Field Images of TCM Grown *C. neoformans* Treated with Glycolysis Inhibitors. a) Representative wider field images of TCM grown wild-type cultures, in the presence and absence of sub-inhibitory concentration of glycolysis inhibitors (2DG or NaCi). Scale bars represent 10 µm. b) Titanides quantification in the presence and absence of sub-inhibitory concentration of glycolysis inhibitors (2DG or NaCi) grown under titan cell inducing conditions. Statistical analysis was performed using unpaired t-test, ns (non-significant). Error bars represent SEM. c) Representative wider field images of India ink staining for capsule visualization of TCM grown wild-type cultures, in the presence and absence of sub-inhibitory concentration of glycolysis inhibitors (2DG or NaCi). Scale bars represent 10 µm. This figure was created using BioRender. https://BioRender.com/3qfzhar.

**Supplementary Figure 2.**
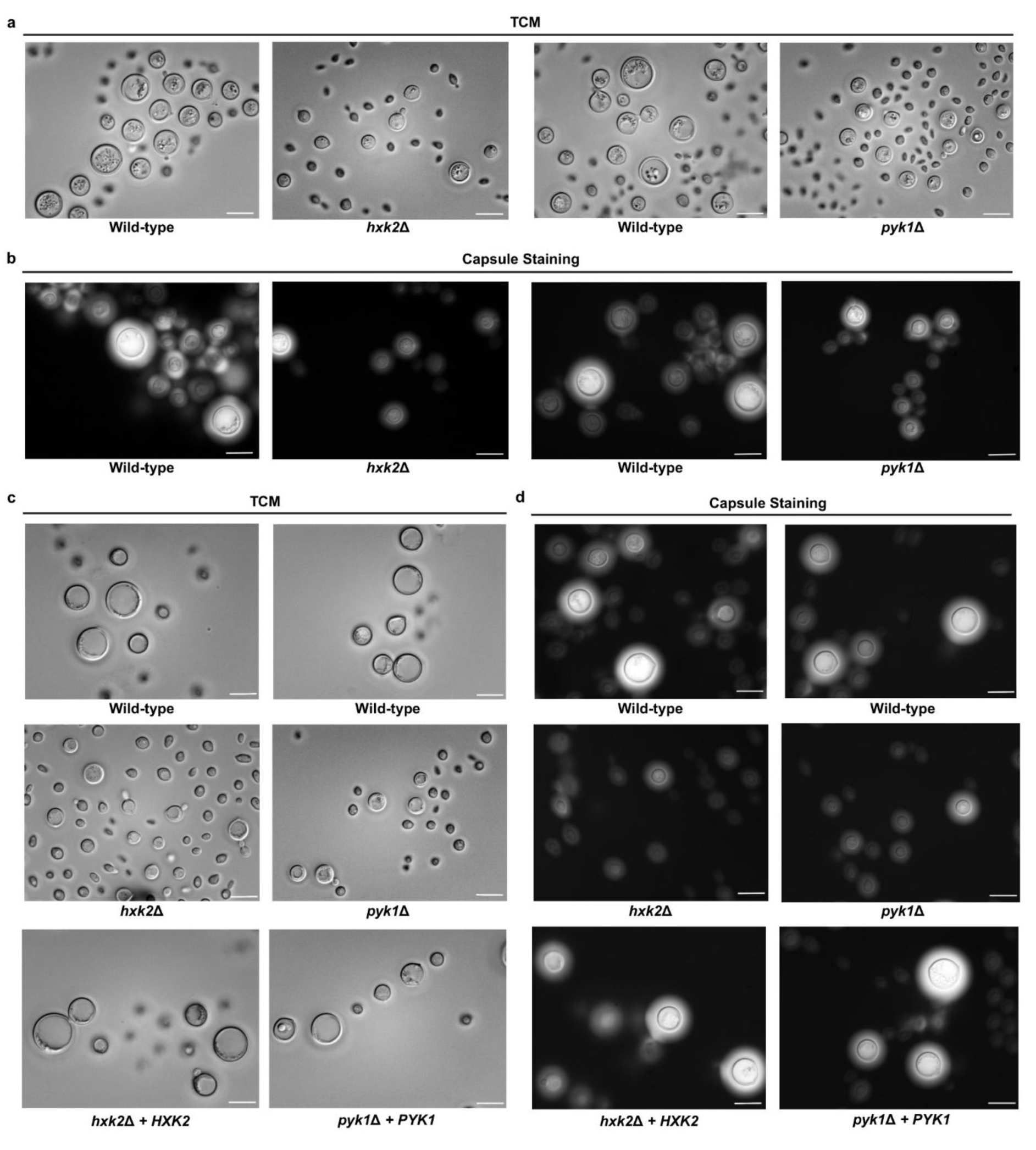
Wider Field Images of TCM Grown Wild-type and Glycolysis Mutants. a) Representative wider field images of wild-type, and glycolysis mutants including *hxk2*Δ, *pyk1*Δ grown under titan cell inducing conditions. Scale bars represent 10 µm. b) Representative wider field images of India ink staining for capsule visualisation of wild-type, and glycolysis mutants including *hxk2*Δ, *pyk1*Δ grown under titan cell inducing conditions. Scale bars represent 10 µm. c) Representative wider field images of wild-type, *hxk2*Δ, *pyk1*Δ, *hxk2*Δ *+ HXK2* and *pyk1*Δ *+ PYK1* strains grown under titan cell inducing conditions. Scale bars represent 10 µm. d) Representative wider field images of India ink staining for capsule visualization of wild-type, *hxk2*Δ, *pyk1*Δ, *hxk2*Δ *+ HXK2* and *pyk1*Δ *+ PYK1* strains grown under titan cell inducing conditions. Scale bars represent 10 µm. This figure was created using BioRender. https://BioRender.com/ivtnjni.

**Supplementary Figure 3.**
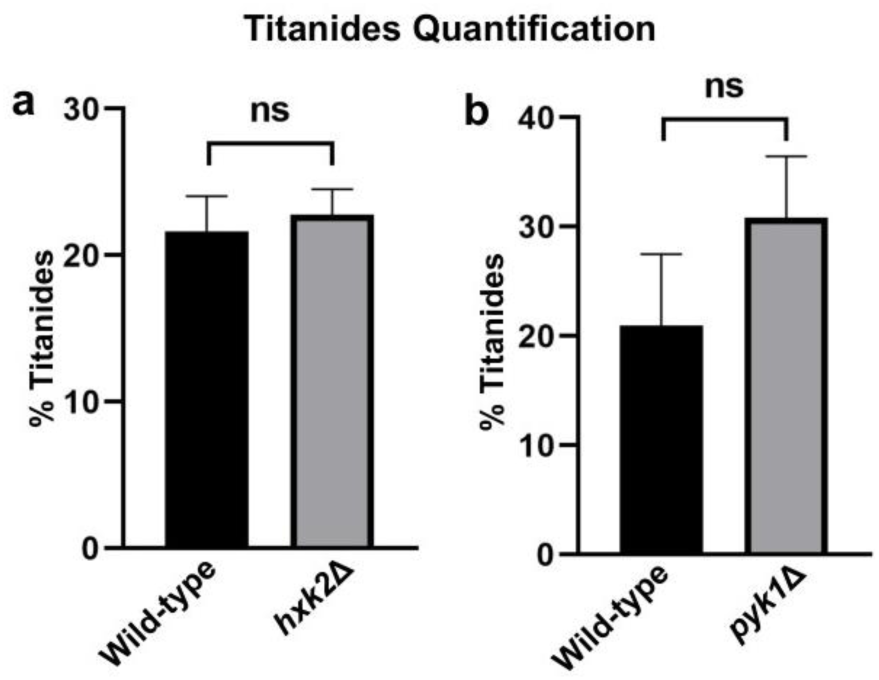
Titanides Quantification in Glycolysis Mutants Grown Under Titan Cell Inducing Conditions. a) Titanides quantification in the wild-type, and glycolysis mutants including *hxk2*Δ grown under titan cell inducing conditions. Statistical analysis was performed using unpaired t-test, ns (non-significant). Error bars represent SEM. b) Titanides quantification in the wild-type, and glycolysis mutants including *pyk1*Δ grown under titan cell inducing conditions. Statistical analysis was performed using unpaired t-test, ns (non-significant). Error bars represent SEM. This figure was created using BioRender. https://BioRender.com/fqdrklv.

**Supplementary Figure 4.**
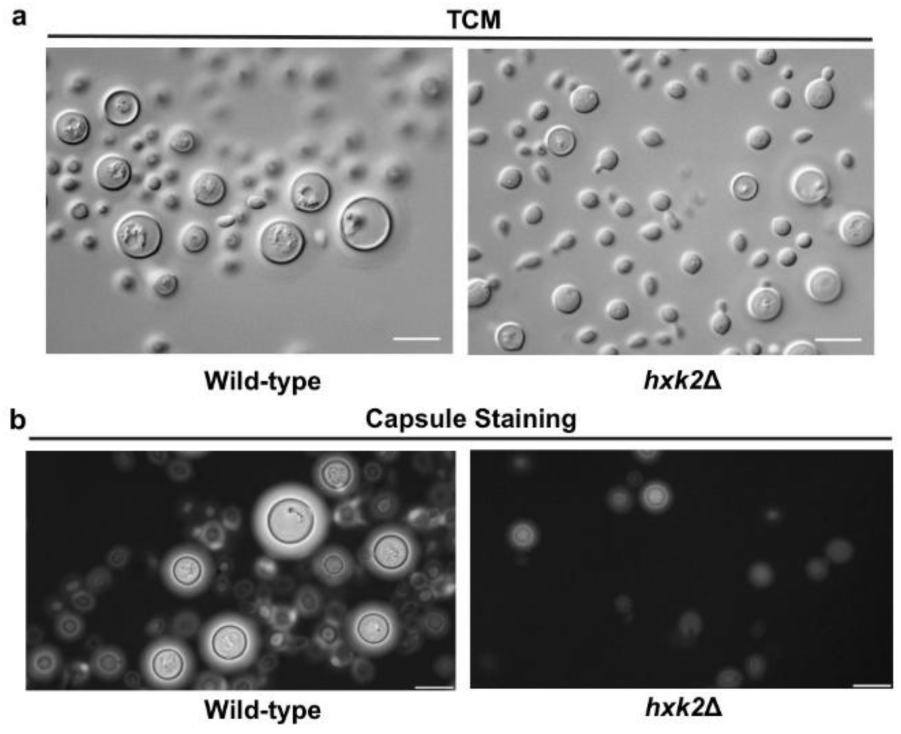
Wider Field Images of 24 Hours TCM Grown Wild-type and Glycolysis Mutants. a) Representative wider field images of wild-type, and glycolysis mutants including *hxk2*Δ grown under titan cell inducing conditions for 24 hours. Scale bars represent 10 µm. b) Representative wider field images of India ink staining for capsule visualization of wild-type, and glycolysis mutants including *hxk2*Δ grown under titan cell inducing for 24 hours. Scale bars represent 10 µm. This figure was created using BioRender. https://BioRender.com/yk6735g.

**Supplementary Figure 5.**
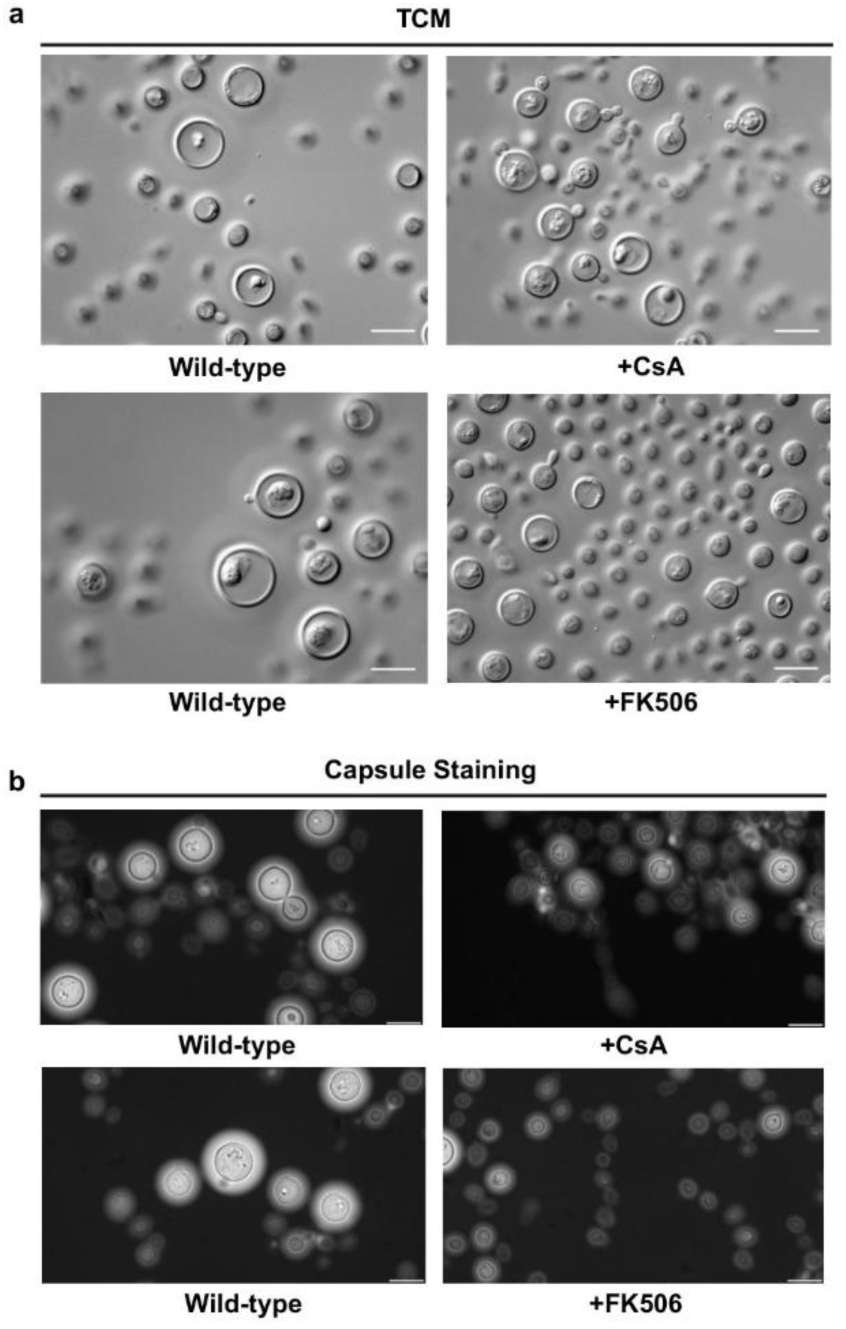
Wider Field Images of TCM Grown *C. neoformans* Treated with Calcineurin Inhibitors. a) Representative wider field images of TCM grown wild-type cultures, in the presence and absence of sub-inhibitory concentration of calcineurin inhibitors (CsA or FK506). Scale bars represent 10 µm. b) Representative wider field images of India ink staining for capsule visualization of TCM grown wild-type cultures, in the presence and absence of sub-inhibitory concentrations of calcineurin inhibitors (CsA or FK506). Scale bars represent 10 µm. This figure was created using BioRender. https://BioRender.com/8rmgwvi.

**Supplementary Figure 6.**
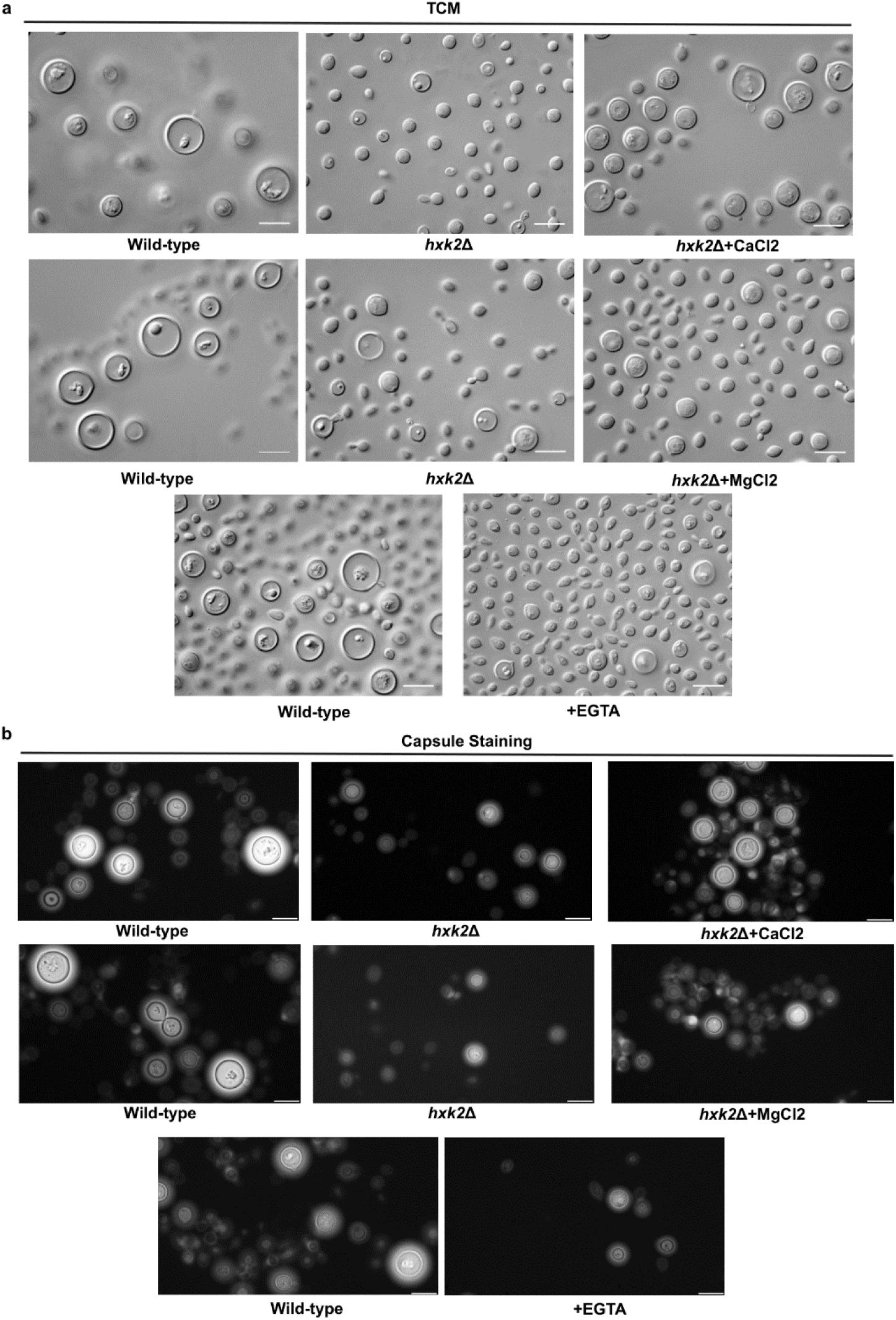
Wider Field Images of TCM Grown *C. neoformans* Treated with CaCl_2_/MgCl_2_/EGTA. a) Representative wider field images of TCM grown wild-type cultures, in the presence and absence of CaCl_2_ (2.5 mM) or MgCl_2_ (2.5 mM) or EGTA (5 mM). Scale bars represent 10 µm. b) Representative wider field images of India ink staining for capsule visualization of TCM grown wild-type cultures, in the presence and absence of CaCl_2_ (2.5 mM) or MgCl_2_ (2.5 mM) or EGTA (5 mM). Scale bars represent 10 µm. This figure was created using BioRender. https://BioRender.com/6sk1ura.

**Supplementary Figure 7.**
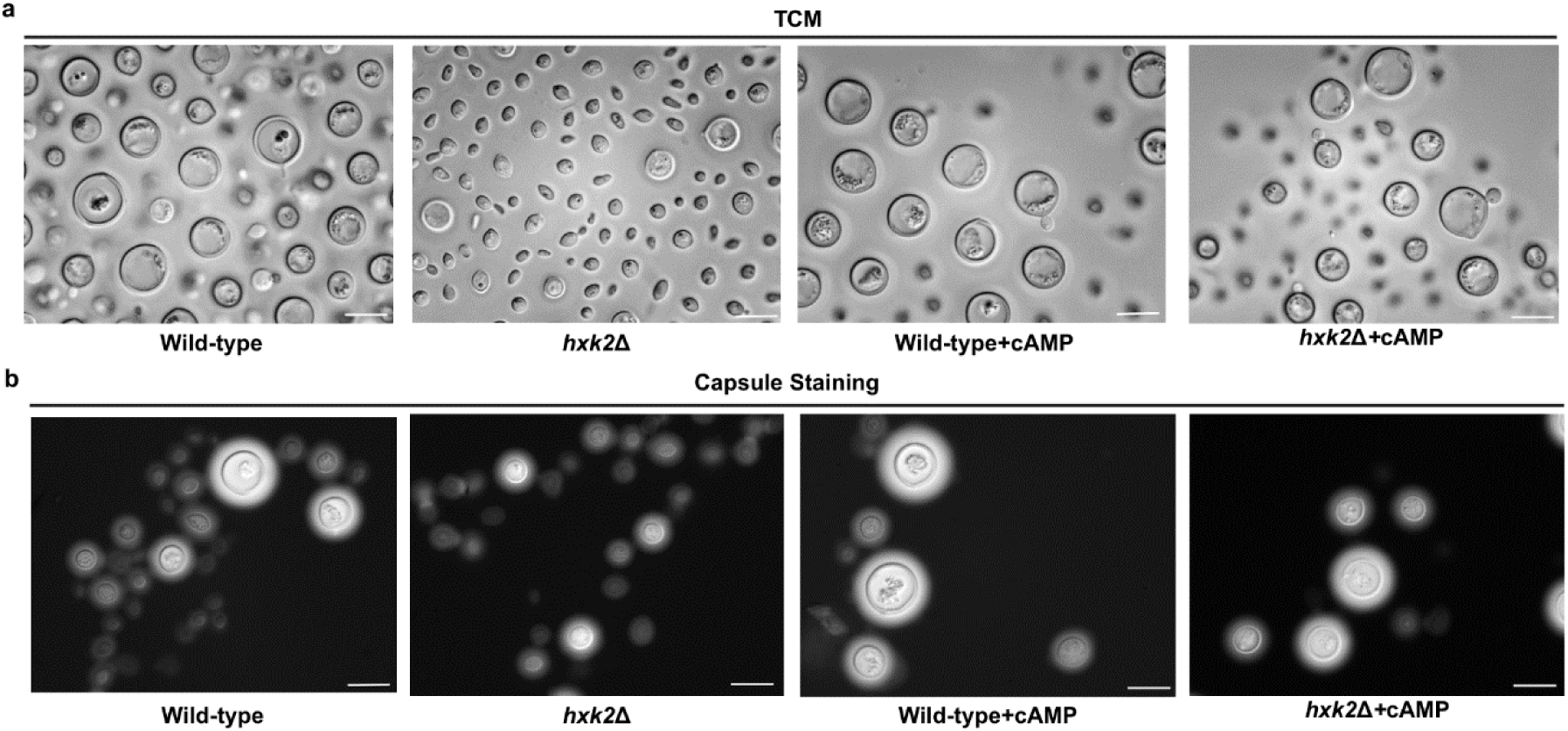
Wider Field Images of TCM Grown *C. neoformans* Exogenously Supplemented with cAMP. a) Representative wider field images of TCM grown wild-type and *hxk2*Δ mutant cultures, in the presence and absence of cAMP (10 mM). Scale bars represent 10 µm. b) Representative wider field images of India ink staining for capsule visualization of TCM grown wild-type *hxk2*Δ mutant cultures, in the presence and absence of cAMP (10 mM). Scale bars represent 10 µm. This figure was created using BioRender. https://BioRender.com/7ruaau4.

